# Dorsal striatal dopamine integrates sensory and reward prediction errors to guide perceptual decisions

**DOI:** 10.64898/2026.01.01.696999

**Authors:** Matthias Fritsche, Chiara Toschi, Yuliia Shevchuk, Olena Didenko, Rathi Puliyadi, Antara Majumdar, Rafal Bogacz, Armin Lak

## Abstract

Perceptual decisions are shaped by expectations about sensory stimuli and rewards, learned through sensory and reward prediction errors. Dopamine is known to convey reward prediction errors that shape perceptual decisions. However, whether dopamine also signals sensory prediction errors during perceptual decision-making remains unknown. We recorded dopamine release in the dorsal striatum of mice performing a visual decision-making task while manipulating sensory and reward expectations. The two manipulations produced similar behavioral biases but elicited opposite dopamine signals, indicating that dopamine within the same striatal region encoded both sensory and reward prediction errors. Sensory prediction error signaling was specific to dopamine as striatal acetylcholine showed no such error signals. Optogenetically stimulating striatal dopamine at stimulus onset biased subsequent perceptual decisions, consistent with updating sensory expectations. A computational model learning from both sensory and reward prediction errors captured behavioral biases and dopamine data, revealing distinct but spatially overlapping dopaminergic teaching signals.

## Introduction

Perceptual decisions rely not only on momentary sensory inputs, but also on sensory and reward expectations shaped by past experience^1–7^. Observers continuously update expectations about upcoming sensory stimuli and rewards based on recent stimulus statistics and decision outcomes, giving rise to robust history-dependent decision biases across species^7–22^. A central question is what neural mechanisms enable the learning and updating of sensory and reward expectations that guide perceptual decisions.

Past work examining how prior sensory expectations are learned converged on the idea that sensory expectations are updated when inputs deviate from predictions, through sensory prediction errors (SPEs). Such SPEs have been prominently observed in cortical neuronal activity^23–30^. In parallel, studies of reward value-guided learning have established that discrepancies between expected and received outcomes, i.e., reward prediction errors (RPEs)^31,32^, are encoded in subcortical systems, particularly in dopamine signals^33–35^. Together, these studies suggest a division of labor in which SPEs arise mainly in cortex and RPEs in dopaminergic and dopamine-recipient subcortical circuits. However, whether dopamine circuits also participate in sensory-predictive learning, and whether sensory- and reward-based teaching signals rely on shared neural substrates remains unknown.

We hypothesized that dopamine signals may convey SPEs as well as RPEs, thereby providing a common teaching signal for multiple forms of learning. Dopamine signals have been extensively studied in reward value-guided reinforcement learning experiments. Classic work shows that midbrain dopamine neurons encode RPEs, with temporal-difference learning models explaining dopamine responses to reward-predicting cues and outcomes^33^. Extensions of these models to partially observable settings also account for more recent findings in perceptual decision-making tasks, where RPEs incorporating statistical decision confidence capture dopamine responses to perceptually uncertain sensory stimuli and rewards^36–42^. However, accumulating evidence suggests that dopamine may have a broader role in sensory processing, beyond RPEs. Midbrain dopamine neurons respond to errors in predicting sensory features of expected rewards^43^, and striatal dopamine release correlates with errors in predicting value-neutral sensory stimuli during associative learning^44,45^. These findings point to the possibility that dopamine signals might encode discrepancies between expected and actual sensory inputs, i.e., SPEs, for updating sensory expectations. However, these earlier studies were not designed to test whether such signals constitute genuine SPEs that influence future perceptual decisions. In parallel, pre-stimulus dopamine responses in the tail of the striatum have been shown to reflect sensory expectations that bias perception^46^. Yet these pre-stimulus dopamine signals represent predictions, rather than SPEs calculated *after* observing the stimulus. Together, these findings point to the involvement of dopamine in sensory processing but leave unresolved whether dopamine conveys SPEs that update sensory expectations during decision-making.

To test whether dopamine signals convey SPEs and RPEs, we trained head-fixed mice on a visual decision-making task. We manipulated the sensory and reward structure of the environment while recording dopamine release in the dorsal striatum - a key region for perceptual decision-making^40,47,48^. Strikingly, manipulating sensory and reward expectations produced similar behavioral biases but elicited opposite dopamine signals. Perceptual decisions were biased towards high-probability expected stimuli and stimuli predicting larger rewards. However, stimulus-evoked dopamine release was *reduced* for high-probability expected stimuli, but *enhanced* for stimuli predicting larger rewards. This opposite scaling with stimulus and reward expectation indicated that dorsal striatal dopamine conveys both SPEs and RPEs. Optogenetic stimulation of stimulus-evoked dopamine signals biased subsequent perceptual choices, supporting the idea that dopamine SPEs update future sensory expectations. Computational modelling further revealed that neither pure reinforcement learning nor sensory-predictive learning models could account for the data; instead, a model that learned jointly from SPEs and RPEs captured both neural and behavioral results. Together, these results indicate that dorsal striatal dopamine conveys overlapping teaching signals that support multiple forms of learning. By encoding both SPEs and RPEs, dopamine extends beyond its traditional reward-centric role to shape perceptual decision-making.

## Results

### Reward and stimulus expectations similarly bias perceptual decisions

We trained mice in a visual decision-making task that included manipulations of reward value or stimulus probabilities (**Fig. 1a**; n = 10 mice, 204 sessions, 71,221 trials). In each trial, we presented a stimulus (grating patch) on the left or right side of a computer screen and mice indicated the stimulus location by steering a wheel with their forepaws, receiving water reward for correct responses. After mice reached expert proficiency on the basic task, we performed two behavioral manipulations. In the reward size manipulation, correct choices to one side yielded a larger reward (3 μL) than the other side (1.5 μL), with the large-reward side switching across blocks of trials (**Fig. 1b**). In the stimulus probability manipulation, reward size was held constant, but in blocks of trials, stimuli were more likely to appear on one side (80/20%; **Fig. 1c**). These manipulations have been widely utilized to probe how learned reward value and sensory priors shape perceptual decisions, showing that these manipulations bias decisions towards more rewarding or more probable alternatives^5–7,49^.

**Fig. 1.**
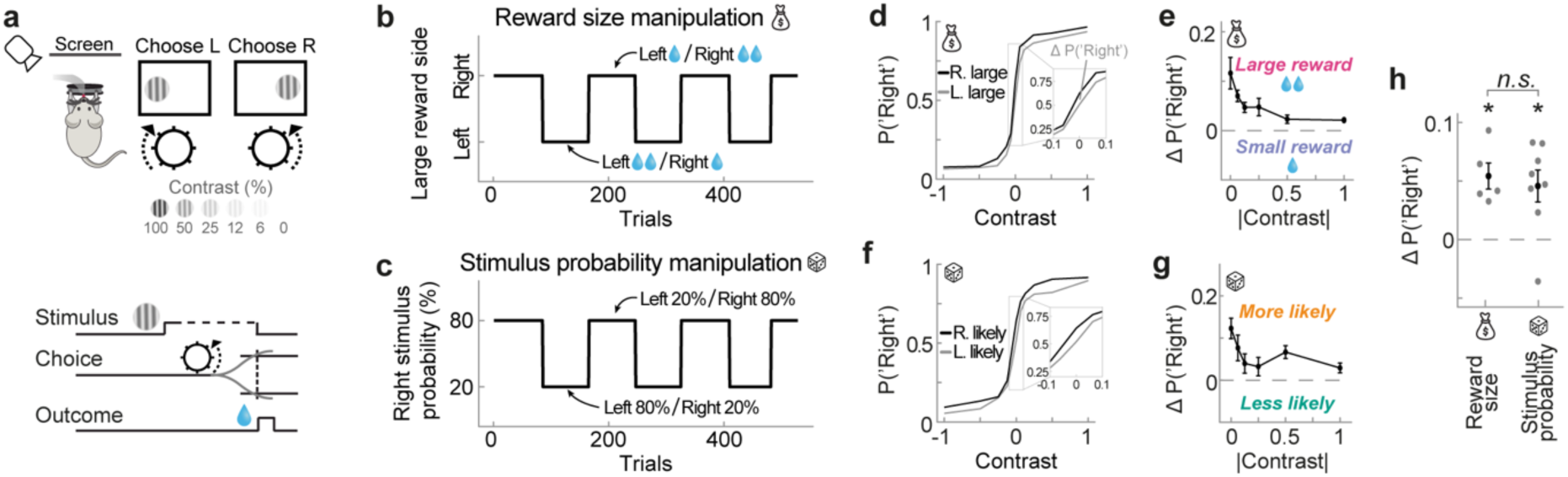
Mice bias visual decisions towards large-reward and high-probability stimuli. **a,** Schematic of the two-alternative visual decision-making task. Head-fixed mice reported the location (left/right) of gratings with varying contrasts by steering a wheel with their forepaws, receiving water reward for correct responses. **b,** In a subset of sessions, reward for correct choices was larger on one side. The large-reward side switched in blocks throughout the session. **c,** In different sessions, stimuli appeared with a higher probability on the left or right side (80/20%), with equal reward for correct choices. **d,** Group-average choices (n = 5 mice, 22,331 trials) in the reward size manipulation, as a function of stimulus contrast (x-axis) and large reward side (black and grey lines). Negative and positive contrasts denote stimuli on the left and right sides, and the y-axis denotes the probability of a rightward choice. Mice exhibit expert performance and bias their choices towards the side with larger reward (see inset for low stimulus contrasts). **e,** Difference between choice probabilities conditioned on the current large reward side (black minus grey lines in panel d, see inset), as a function of absolute stimulus contrast. Positive y values indicate a tendency to choose the large reward side. Data points show group averages. Error bars in all panels denote SEMs. **f,** Group-average choices (n = 8 mice, 48,890 trials) in the stimulus probability manipulation. Same as in d, but conditioned on the higher probability side. Mice bias their choices towards the more likely side (see inset for low stimulus contrasts). **g,** Difference between choice probabilities conditioned on the current high probability side (black minus grey lines in panel f), as a function of absolute stimulus contrast. Positive y values indicate a tendency to choose the more likely side. **h,** Choice bias towards the larger reward and higher probability sides, averaged across all stimulus contrasts. Mice exhibited choice biases of similar magnitude across the two manipulations. *p<0.05. See also Fig. S1.

Mice showed robust decision biases, manifested as a shift in their psychometric curves while maintaining high sensitivity to visual stimuli (**Fig. 1d,f**), consistent with prior work. They preferentially chose the larger-reward side (**Fig. 1d-e**; mean bias 5.4 ± 1.1%, t(4) = 4.87, p = 0.008) and the higher-probability side (**Fig. 1f-g**; mean bias 4.6 ± 1.4%, t(7) = 3.36, p = 0.01). Both biases were largest in trials in which the presented stimulus contrast was low, suggesting that mice particularly relied on reward and probability context when they were perceptually uncertain. Importantly, the two manipulations produced biases of comparable magnitude (**Fig. 1h**; t(10.93) = 0.48, p = 0.64), allowing us to compare their neural correlates.

Response times depended on stimulus contrast (**Fig. S1a-b**; reward size manipulation: F(1,4) = 23.89, p = 0.008; stimulus probability manipulation: F(1.14,7.96) = 15.64, p = 0.004, repeated-measures ANOVAs), but did not significantly depend on stimulus probability or impending reward size (**Fig. S1c–e**). Pupil responses, measured through videography, revealed a selective effect of sensory expectation: surprising stimuli evoked larger post-stimulus pupil dilations than expected stimuli (**Fig. S1f-i**; t(8) = 3.54, p = 0.009), consistent with previous reports linking pupil size to sensory prediction errors^50,51^. In contrast, reward asymmetry did not significantly affect pupil size (**Fig. S1j**; t(4) = 0.50, p = 0.64).

Together, these results show that mice flexibly adapt their perceptual choices in line with reward and stimulus expectations, producing similar decision biases in each case. Computational theories attribute these adaptations to prediction error-based learning^22,50,52^. Specifically, RPEs reinforce choices that yield unexpectedly large rewards, while SPEs adjust sensory expectations when sensory events deviate from prior predictions. We therefore next asked whether dorsal striatal dopamine encodes such prediction errors during perceptual decision-making.

### Reward size and stimulus probability oppositely modulate striatal dopamine

We measured dopamine release in the dorsolateral striatum (DLS), using dopamine sensors^53^ (GRAB_DA2m_) and fiber photometry while mice performed our visual decision-making tasks (**Fig. 2a-b**). Dopamine release showed phasic increases at distinct times in the trial (**Fig. 2c**). A linear encoding model revealed that these dopamine responses were mostly driven by stimuli and rewards (6 and 12% unique explained variance, see Methods), with negligible contributions from wheel movements to report a choice (**Fig. 2d**; 0.4%), consistent with previous studies^39,40^.

**Fig. 2.**
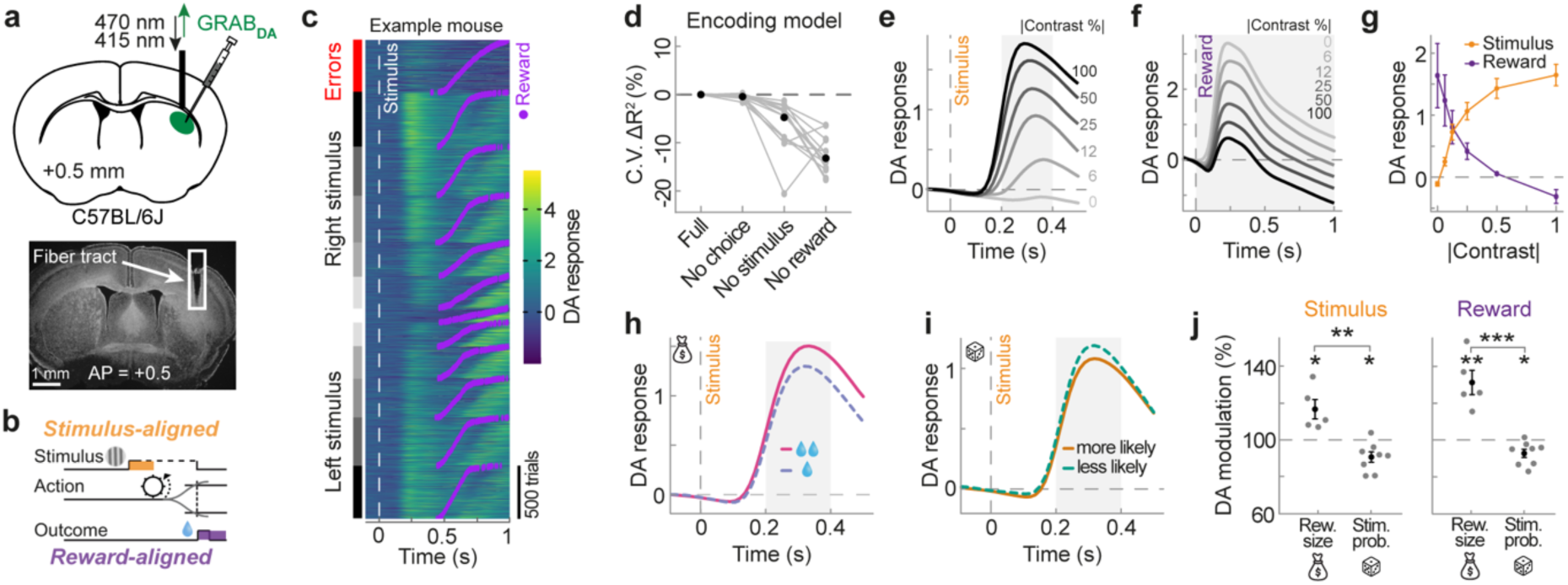
Striatal dopamine increases for large-reward stimuli, but decreases for high-probability stimuli, reflecting RPEs and SPEs. **a,** Top, Schematic of fiber photometry in the dorsolateral striatum (DLS), imaging dopamine release using GRAB_DA2m_. Bottom, Example of post-mortem verification of fiber placement. **b,** Schematic of stimulus-aligned (yellow) and reward-aligned (purple) analysis time windows (see panels e-j). **c,** Trial-by-trial dopamine responses of an example animal, aligned to stimulus onset (white dashed line) and sorted by trial type (left column) and outcome time (purple dots). **d,** Linear encoding model of dopamine release: Cross-validated explained variance was most reduced when removing reward or stimulus regressors (see Methods; data pooled across manipulations). Grey lines denote individual animals, black dots denote group averages. **e,** Group-average dopamine response aligned to stimulus onset (gray dashed line), split by absolute stimulus contrast (gray to black; correct trials only). Gray shaded area indicates the stimulus time period over which we averaged stimulus responses (g-j; excluding time points after reward delivery). See Fig. S2 for separate responses in reward size and stimulus probability manipulations. **f,** Same as in e, but aligned to reward onset. **g,** Group-average stimulus-evoked (yellow) and reward-evoked (purple) dopamine responses as a function of absolute stimulus contrast (rewarded trials only; averaged over gray shaded areas in e and f). Error bars denote SEMs. **h,** Stimulus-evoked dopamine release was increased when the stimulus was presented at the large (solid pink) versus small reward side (dashed purple). Response averaged across all contrast levels and animals. See Figure S2 for responses per contrast level. **i,** Stimulus-evoked dopamine release was decreased when the stimulus was presented at the high (solid orange) versus low probability side (dashed green). **j,** Modulation of dopamine release by reward size and stimulus probability manipulations at stimulus (left) and reward time (right). Modulation was computed by dividing high by low reward/probability responses. Reward modulation was computed for trials without visual stimuli (0% contrast), and conditioned on high/low reward/probability choices, to avoid overlapping stimulus-evoked responses. Grey dots denote individual animals, black dots show group means. *p<0.05, **p<0.01, ***p<0.001. See also Fig. S2.

Dopamine responses to stimuli and rewards scaled with stimulus contrasts in both behavioral tasks. Stimulus onset evoked dopamine responses that scaled positively with stimulus contrast (**Fig. 2e**; F(1.31,9.19) = 68.38, p = 7.83 x 10-6 ). Conversely, reward-evoked dopamine responses scaled negatively with contrast, peaking when rewards followed trials without visual stimuli, i.e. 0% contrast trials (**Fig. 2f**; F(1.54, 10.75) = 111.03, p = 1.42 x 10-7). Thus, stimulus and reward responses were inversely related to sensory evidence (**Fig. 2g**). These patterns held for both manipulations separately (**Fig. S2a-c & k-m**) and were independent of stimulus side (**Fig. S2d,n**). This inverse scaling is consistent with RPE-coding of temporal difference learning models: dopamine encodes the expected reward value during stimulus processing, with high contrast stimuli predicting a highly certain reward and it encodes RPE during outcome, where maximal surprise occurs when receiving a reward given a maximally uncertain stimulus. At the same time, the positive scaling of stimulus-evoked dopamine with stimulus contrast also fits with SPE signaling, where more intense or informative sensory events produce greater deviations from prior sensory expectations. Thus, while the results are fully compatible with RPE signaling, they are also consistent with SPEs, making it difficult to distinguish the two based on these results alone.

Stimulus-evoked dopamine responses were oppositely modulated by reward size and stimulus probability, indicating that dopamine signals encode both RPEs and SPEs. Stimulus-evoked dopamine scaled *positively* with reward size. Large-reward stimuli evoked stronger dopamine responses than small-reward stimuli (116.7 ± 5.3% relative to small reward, t(4) = 3.44, p = 0.026; **Fig. 2h,j**). This matches RPE signaling: Large-reward visual cues set higher reward expectations. In contrast, dopamine scaled *negatively* with stimulus probability: High-probability stimuli evoked weaker dopamine responses than low-probability stimuli (90.6 ± 2.8% relative to low probability, t(7) = -3.30, p = 0.013; **Fig. 2i,j**). Thus, dopamine signals showed opposite modulations by pending reward size and stimulus probability, despite similar behavioral biases (t(4.71) = 7.90, p = 0.002). Crucially, the decreased dopamine response to high-probability stimuli cannot be explained by RPEs. This is because high-probability stimuli are more frequently rewarded and therefore have a higher expected value. Consequently, RPE coding predicts increased, not decreased dopamine responses to these stimuli (see computational modeling below). Instead, the result is consistent with SPE signaling, where the presentation of a low-probability stimulus violates prior sensory expectations, and evokes a larger SPE. Together, these results suggest that alongside RPEs, dopamine responses signal SPEs.

Further analyses ruled out alternative explanations for these dopamine modulations, based on confounding choice-related signals. First, response times were not significantly affected by stimulus value or probability (**Fig. S1c-e**). Second, the encoding model showed negligible contributions from choice on dopamine signals (<1%; **Fig. 2d**). Consistently, dopamine peak latencies showed a weak negative, not positive, relationship with response times (**Fig. S2e-f,o-p**; reward size: t(4) = –2.69, p = 0.055; probability: t(7) = –3.18, p = 0.015). Third, no modulation was observed in stimulus-absent trials (i.e. zero-contrast trials), when conditioning on low/high reward/probability choices (**Fig. S2i,s**), and qualitatively similar modulations were observed on trials with incorrect responses (**Fig. S2j,t**). Finally, opposite modulations also remained on trials with responses outside of our stimulus analysis window (RT > 0.4s; reward size: t(4) = -3.59, p = 0.023; stimulus probability: t(7) = 4.69, p = 0.002). Together, these controls link the modulation to stimulus processing rather than motor behavior.

Both reward size and stimulus probability modulated dopamine responses also at the time of reward delivery. As expected, reward-evoked dopamine was relatively increased for large compared to small water rewards (**Fig. 2j**, right; 131.3% +- 6.6% of small reward response, t(4) = 5.43, p = 0.006). In contrast, dopamine was decreased when receiving reward following a high rather than low probability stimulus (92.7% +- 2.2% of low probability response; t(7) = -3.16, p = 0.016). Therefore, similar to stimulus responses, reward-evoked dopamine was oppositely modulated by reward size and stimulus probability (**Fig. 2j**; t(6.27) = 6.01, p = 0.0008). This opposite modulation at reward time is consistent with RPE signaling, as larger water rewards lead to increased RPEs, but rewards following high- probability, and therefore frequently rewarded stimuli, are more expected and therefore lead to decreased RPEs (see also computational modeling section below).

In summary, despite similar behavioral choice biases in response to asymmetric rewards and stimulus priors, dorsal striatal dopamine responses to stimuli exhibited opposite modulations, suggesting that they encode overlapping RPEs and SPEs.

### Dopaminergic SPEs occur in absence of instructed actions

We proposed above that enhanced dopamine responses to low-probability stimuli reflect SPEs. An alternative possibility is that they reflect action prediction errors that arise when a rare stimulus drives a less frequently taken action^54–56^. Although our results so far linked dopamine responses to stimulus processing and not action execution, we designed a new experiment to conclusively distinguish between these possibilities.

We trained a separate cohort of mice in a visual classical conditioning task (**Fig. 3a**; n = 6 mice, 64 sessions, 11,675 trials). On each trial, mice viewed a one-second grating on the left or right side of a screen. Each stimulus was followed by a fixed water reward (3 μL), with inter-trial intervals of 6 to 60s. We manipulated sensory expectations by biasing the probability of stimuli appearing on the left or right side across experimental sessions (80/20% probability), similar to the decision-making task described above. Importantly, mice did not have access to the response wheel, and rewards were not contingent on behavior. Thus, sensory expectations were manipulated in the absence of instructed actions. We measured dorsal striatal dopamine release, while mice passively experienced the stimuli and reward.

**Fig. 3.**
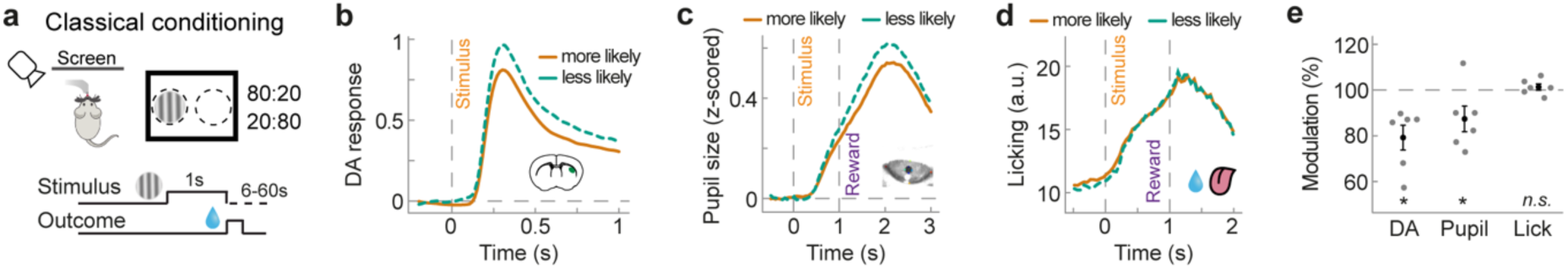
Dopaminergic SPEs occur in the absence of instructed actions. **a,** Schematic of the visual classical conditioning paradigm. Head-fixed mice were presented with 100% contrast gratings, appearing on the left or right side. All grating presentations were followed by a water reward after 1 second. On a given day, gratings were more likely to appear on the left or right side (80/20% probability). **b,** Group-average dopamine response (n = 6 mice, 11,675 trials) with respect to stimulus onset (gray dashed line), split by whether the stimulus appeared on the more (solid orange) or less likely side (dashed green). For reward response see Fig. S3. **c,** Group-average pupil size with respect to stimulus onset. **d,** Group-average licking activity with respect to stimulus onset (see Methods). **e,** Modulation of stimulus-evoked dopamine, pupil size and pre-reward, anticipatory licking by stimulus probability. Modulation was computed by dividing the response to high probability, surprising stimuli, by that of low probability, expected stimuli. DA stimulus and anticipatory licking time window: 0-1s post stimulus; Pupil size time window: 1-3s post stimulus. Grey dots denote individual animals, black dots show group means, error bars denote SEMs. *p<0.05. See also Fig. S3.

Dorsal striatal dopamine signals reflected sensory expectations in the classical conditioning task, similar to the decision-making task. High-probability, expected stimuli evoked weaker dopamine release than low-probability stimuli (**Fig. 3b,e**; 79.2 ± 5.4% of low probability response, t(5) = -3.31, p = 0.011). Low-probability, surprising stimuli also produced larger pupil dilations (**Fig. 3c,e**; 87.4 ± 5.6%, t(5) = -2.36, p = 0.032), replicating results from the instrumental task (see **Fig. 2i** and **Fig. S2i**). In contrast, anticipatory pre-reward licking did not differ between low- and high-probability stimuli (**Fig. 3d,e**; 101.3 ± 1.4%, t(5) = 0.94, p = 0.39).

Together, these results show that sensory expectations modulate stimulus-evoked dorsal striatal dopamine even in the absence of instructed actions, supporting the encoding of sensory- rather than action-prediction error.

### Striatal acetylcholine release does not encode SPEs

We next asked whether encoding of SPEs is specific to dopamine, or reflects a more general feature of striatal neuromodulation. The striatum contains high levels of acetylcholine, released by local cholinergic interneurons^57,58^. Cholinergic interneurons respond to salient sensory events^59–61^, and striatal acetylcholine has been implicated in driving local dopamine release^62^. Thus, striatal acetylcholine might itself contribute to dopaminergic SPEs. We therefore measured DLS acetylcholine release using GRAB ^63^ during the visual decision task (**Fig. 4a**; n = 11 mice, 152 sessions, 58,651 trials). Acetylcholine signals were clearly modulated during trials (**Fig. 4b**). A linear encoding model attributed most unique variance to stimulus and reward events (15% and 24%), with smaller choice contributions (9%; **Fig. S4b**).

**Fig. 4.**
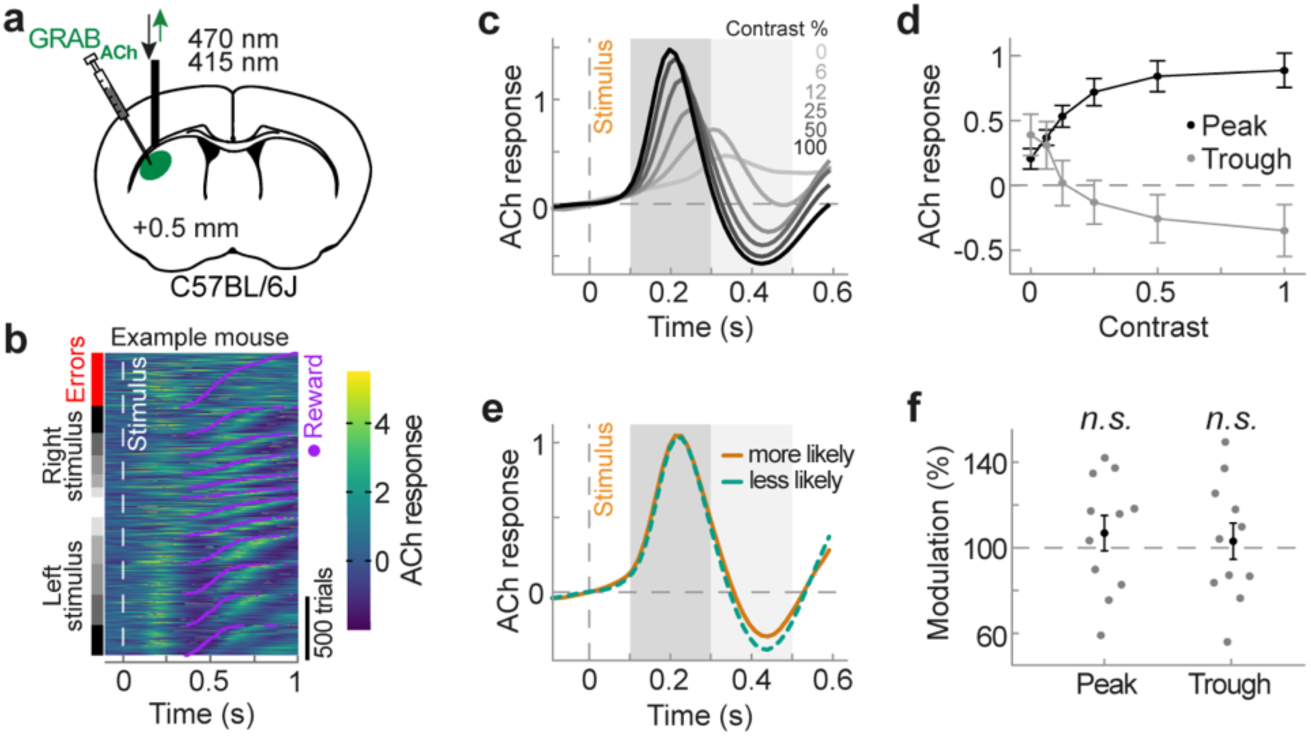
Striatal acetylcholine release does not encode SPEs. **a,** Schematic of fiber photometry in the DLS, imaging acetylcholine release using GRAB_ACh3.0_. Acetylcholine release was recorded from the opposite hemisphere of the same animals in which dopamine release was measured, as well as in a small number of additional animals (total n = 11 mice, 58,651 trials, see Methods). **b,** Trial-by-trial acetylcholine responses of an example animal, aligned to stimulus onset (white dashed line) and sorted by trial type (left column) and outcome time (purple dots). **c,** Group-average acetylcholine response aligned to stimulus onset (gray dashed line), split by absolute stimulus contrast (gray to black; correct trials only). Dark and light gray shaded areas indicate the time periods over which we averaged peak and trough stimulus responses (d-f; excluding time points after reward delivery). See Fig. S4 for reward responses. **d,** Average peak (black) and trough (grey) acetylcholine stimulus responses as a function of contralateral stimulus contrast (rewarded trials only; averaged over gray shaded areas c). Data points show group means and error bars denote SEMs. **e,** Group-average stimulus-evoked acetylcholine release, split by whether the stimulus appeared on the more (solid orange) or less likely side (dashed green). Responses were averaged across all contralateral contrast levels. **f,** Modulation of peak (left) and trough (right) acetylcholine response by stimulus probability manipulation. Modulation was computed by dividing responses to high-probability by low-probability stimuli. Grey dots denote individual animals, black dots show group means. Error bars denote SEMs. See also Fig. S4.

Striatal acetylcholine responses to stimuli did not encode SPEs. Stimulus onset evoked a biphasic acetylcholine response with an early peak (∼200 ms) followed by a trough (**Fig. 4c**), consistent with previous studies^64,65^. Both peak and trough scaled with visual contrast (**Fig. 4d**, peak: F(1.60,15.98) = 29.95, p = 9x10-6; trough: F(1.08, 10.83) = 8.84, p = 0.012). Unlike dopamine, acetylcholine responses were strongly lateralized to contralateral stimuli (**Fig. S4c**, main effect of stimulus side, peak: F(1,10) = 42.24, p = 6.9x10-5; trough: F(1,10) = 1.37, p = 0.27; contrast x side interaction, peak: F(4,40) = 14.18, p = 2.7x10-7; trough: F(1.37, 13.67) = 4.57, p = 0.041). We therefore focused on contralateral trials, though results held when including all trials. High-probability stimuli produced peak and trough responses comparable to low-probability stimuli (**Fig. 4e-f**, peak: 106.8% +- 8.2%, t(10) = 0.39, p = 0.71; trough: 103.1% +- 8.5%, t(10) = -0.60, p = 0.95). This result occurred, despite animals showing clear probability-driven choice biases (**Fig. S4j-k**, 6.1% ± 1.3%, t(10) = 4.63, p = 0.0009). Reward-evoked acetylcholine responses were likewise not significantly modulated by stimulus probability (**Fig. S4d,l,m**).

In sum, striatal acetylcholine robustly encoded lateralized visual information, but not SPEs. This specificity also highlights dopamine as the carrier of SPE signals in dorsal striatum.

### Dopaminergic SPEs boost future perceptual choice biases

We next asked whether DLS dopamine signals contribute to updating sensory priors that guide future perceptual decisions. Dopamine is well known to drive learning as demonstrated by causal manipulations that mimic outcome-evoked dopamine RPEs. Yet, it remains unclear whether stimulus-evoked dopamine SPEs can influence subsequent perceptual decisions. To test this, we used optogenetic stimulation to mimic dopaminergic SPEs during stimulus presentation and examined their impact on future choices. We expressed the excitatory opsin FLEX-ChrimsonR in dopamine neurons of DAT-Cre mice by injecting the virus into the substantia nigra pars compacta (SNc). We then optogenetically stimulated DLS dopamine release by delivering brief laser pulses through an implanted optic fiber (**Fig. 5a-b**), while mice performed the visual decision task in the in the absence of reward size or stimulus probability manipulations (**Fig. 5c**; n = 10 mice, 90 sessions, 32,148 trials). On 40% of randomly interleaved trials, we delivered brief laser stimulation (250 ms) at stimulus onset. We then analyzed behavior on subsequent, non-stimulated trials, thus examining the learning effects of these signals in trials without any laser stimulation (**Fig. 5d**).

**Fig. 5.**
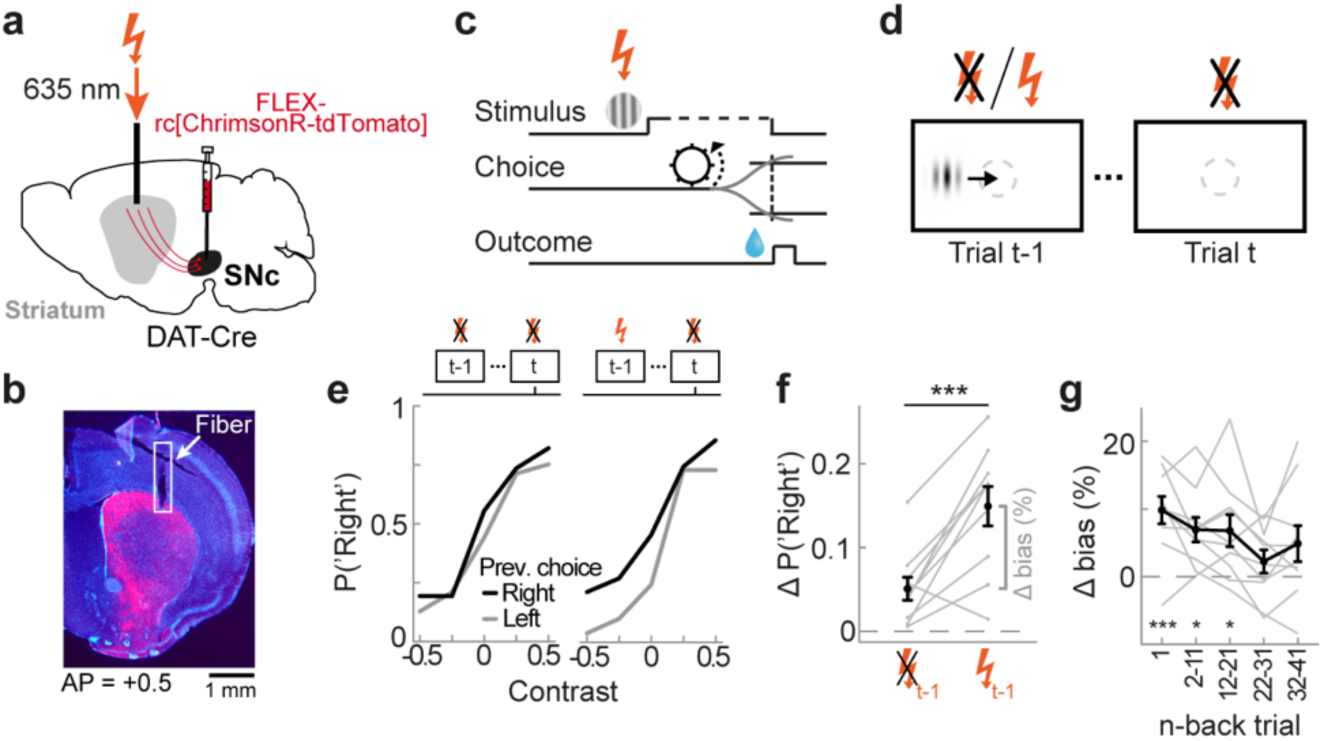
Optogenetic dopamine stimulation at stimulus onset increases subsequent perceptual decision biases. **a,** Schematic of the optogenetic stimulation experiment. We injected FLEX-ChrimsonR in the SNc of DAT-Cre mice and stimulated dopamine axons in the DLS through an optic fiber, while mice performed the decision-making task (n = 10 mice, 32,148 trials). **b,** Example of post-mortem verification of fiber placement and virus expression. **c,** We delivered light stimulation for 250 ms following stimulus onset. **d,** Choice behavior on non-stimulation trials (t) was analyzed as a function of whether the preceding rewarded high-contrast trial (t-1) was paired with stimulation. **e,** Group-average choice behavior on trial t as a function of current stimulus contrast (x-axis) and previous choice (black and grey lines), separately for previously stimulated (left) or non-stimulated trials (right). **f,** Difference between choice probabilities conditioned on the previous choice (black minus grey lines in panel e), averaged across all contrast levels. Positive y values denote a tendency to report the same side as the previous stimulus. Following stimulated trials, animals show a higher tendency to report the same side as the previous stimulus. Grey dots denote individual animals, black dots show group means. Error bars denote SEMs. **g,** The average increase in stimulation-driven choice repetition is largest when stimulating on the previous trial (1-back), but remains statistically significant when stimulating 2-11 and 12-21 trials back. Grey dots denote individual animals, black dots show group means. Error bars denote SEMs. *p<0.05, **p<0.01, ***p<0.001. See also Fig. S5.

Optogenetic dopamine stimulation increased choice repetition after rewarded, high-contrast trials. Mice were more likely to choose the same stimulus alternative on the next trial (**Fig. 5e-f**, t(9) = -4.87, p = 0.0009, previous correct trials only). This effect was not explained by reinforcing actions. It persisted even when the preceding choice occurred >1 s after laser offset (**Fig. S5f**, t(9) = -2.61, p = 0.028). Moreover, it was absent when stimulation occurred on zero-contrast trials without a visual cue (t(9) = 0.30, p = 0.77). The optogenetically-induced bias was long-lasting, consistent with dopamine-dependent plasticity. Stimulation continued to increase repetition biases when delivered up to 12-21 trials back (**Fig. 5g**, t(9) = 2.89, p = 0.03, Bonferroni-corrected for multiple comparisons). Finally, stimulation also produced immediate effects: faster responses on zero-contrast trials and lower choice accuracy (**Fig. S5b-c**). However, these immediate changes did not correlate with the subsequent choice bias (**Fig. S5d-e**).

Together, these results show that dopamine release at stimulus onset induces future perceptual choice biases, supporting a causal role for dopaminergic SPEs in updating sensory priors.

### Concurrent perceptual inference and reinforcement learning explain opposite dopamine modulation

To formally test whether dopamine signals encode SPEs, in addition to RPEs, and how these signals shape behavior, we next built computational models of agents performing the same visual decision-making tasks as our mice and asked which learning framework best accounted for both the behavioral and dopamine data.

A model which only performed RPE-guided reinforcement learning (RL) could not account for the dopamine signals. We first implemented a RL agent that learned to associate noisy sensory stimuli with choices to maximize reward^66^ (**Fig. 6a**, blue box; see Methods). In this framework, dopamine reports a temporal difference RPE (**Fig. 6b**; TD-RPE). When fit to the behavioral data, this agent reproduced key aspects of the mice’s performance: it developed decision biases toward more highly rewarded and more probable stimuli, reflecting reinforcement of actions that yielded larger or more frequent rewards (**Fig. S6d**). It also captured the inverse dependence of stimulus- and reward-evoked dopamine on visual contrast, consistent with a confidence-weighted TD-RPE (**Fig. S6d**). However, it failed to reproduce opposite dopamine modulation by reward size and stimulus probability at stimulus onset. Instead, it predicted stronger dopamine responses for both large-reward and high-probability stimuli (**Fig. S6d**). Thus, pure RL could not account for the stimulus-evoked dopamine signals.

**Fig. 6.**
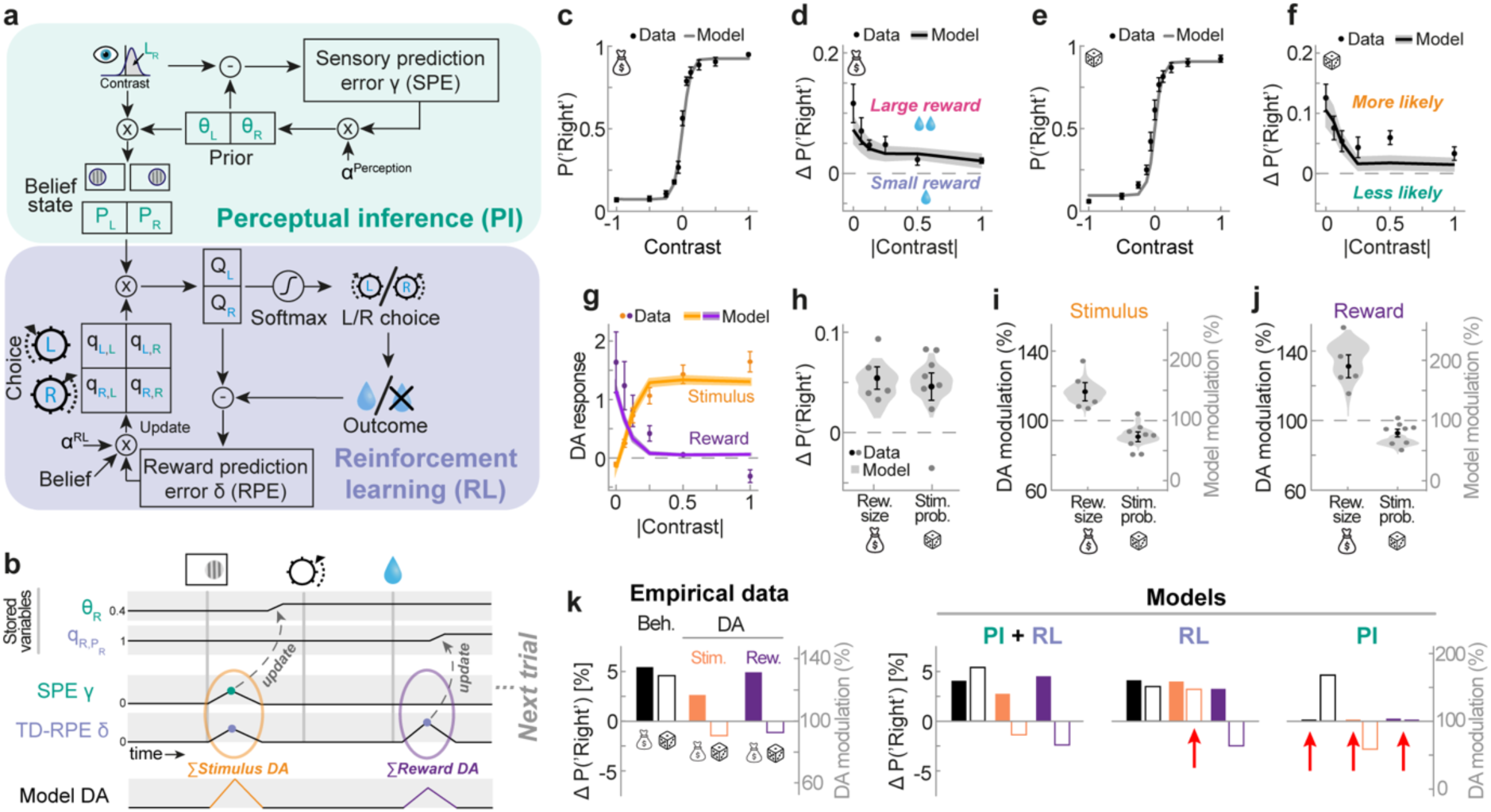
Bayesian perceptual inference and reinforcement learning jointly explain similar behavioral biases but opposite dopamine modulations. **a,** Schematic of reinforcement learning (blue) with Bayesian perceptual inference (green; PI+RL model, see Methods). **b,** Schematic of within-trial changes of model variables (top four rows) and putative dopamine signals (bottom row). **c,** Psychometric curves PI+RL model (lines) and mice (points) in the reward size manipulation. The model accurately captures the mice’s dependence of choice (y-axis) on current contrast (x-axis). **d,** Difference between choice probabilities conditioned on the current large-reward side (black minus grey lines in panel c) for PI+RL model (line) and mice (points). Shaded error regions denote s.d. across 100 bootstrapped model fits (see Methods). **e,f,** Same as panels c and d for stimulus probability manipulation. **g,** PI+RL model stimulus-evoked (yellow) and reward-evoked (purple) dopamine responses as a function of absolute stimulus contrast (rewarded trials only). Model dopamine was taken as a weighted sum of SPEs and temporal-difference RPEs (see panel b). Shaded error regions denote s.d. across 100 bootstrapped model fits. **h,** Choice bias towards the lager-reward and higher-probability sides, averaged across all stimulus contrasts, for PI+RL model (violin plots, 100 bootstrapped model fits) and mice (data points). **i,** Dopamine modulation by reward size and stimulus probability at time of stimulus for PI+RL model (violins) and mice (data points). **j,** same as in i but at time of reward. **k,** Left, summary of empirical behavioral bias and dopamine patterns across reward size and stimulus probability manipulations. Right, Only the PI+RL model recapitulates similar behavioral biases, but opposite dopamine modulations. Red arrows highlight mismatches between model predictions and empirical data. See also Fig. S6.

A model that learned from both RPEs and SPEs accounted for the dopamine data. We next extended the model with a Bayesian perceptual inference (PI) process. This agent infers the stimulus location by combining noisy sensory evidence with a prior expectation derived from recent stimulus history (**Fig. 6a**, green box; see Methods). The prior is updated by SPEs - mismatches between expected and observed sensory inputs. In addition to RPEs, dopamine was allowed to signal an unsigned SPE, and total dopamine activity was modeled as a weighted mixture of the two (**Fig. 6b** and Methods). This combined agent captured the behavioral data better than either process alone (RL: ΔBIC = 304, D(2) = 327, p < 10-16, likelihood ratio test; PI: ΔBIC = 69, D(2) = 92, p < 10-16). Behaviorally, it reproduced psychometric shifts toward both large-reward and high-probability stimuli (**Fig. 6c-f,h**). The model’s dopamine signals also mirrored empirical recordings: stimulus- and reward-evoked dopamine showed opposite dependencies on visual contrast (**Fig. 6g**), and dopamine was oppositely modulated by reward size and stimulus probability, both at stimulus and reward times (**Fig. 6i-j**). These opposite modulations emerged naturally from the mixture of SPEs and RPEs, but neither signal alone could generate them (**Fig. S6b-c**).

In sum, we show that previously proposed frameworks of RL and PI are each insufficient in explaining our results (**Fig. 6k**). Instead, a model that combines both types of learning accounted for the results, indicating that mice simultaneously engage in PI and RL and that DLS dopamine reflects overlapping SPEs and RPEs. Crucially, while both forms of learning are simultaneously active, differences in stimulus and reward statistics in a given context can differentially engage PI and RL mechanisms, leading to similar behavioral decision biases, but opposite dopamine modulations.

## Discussion

Manipulating sensory and reward expectations in a visual decision task produced similar biases in perceptual decisions, yet elicited opposite patterns of dopamine release. This dissociation revealed overlapping SPEs and RPEs in dorsal striatum dopamine signals. Stimulus-evoked dopamine increased when visual stimuli violated sensory expectations, even in the absence of instructed actions. This modulation was specific to dopamine, as acetylcholine showed no such error signals. Optogenetically stimulating dopamine release at stimulus onset biased subsequent perceptual decisions, consistent with updating sensory expectations. Finally, computational modeling showed that both perceptual decision-making behavior and dopamine dynamics were best explained by a framework combining perceptual inference and reinforcement learning, in which observers learn from both SPEs and RPEs.

Dopamine is widely recognized to encode RPEs, guiding learning from positive and negative outcomes during reward value-guided decision-making^33,34,67^. Recent studies have shown that dopaminergic RPEs also occur during perceptual decision-making^36,37,39,41,42^: extending a TD-RL model to incorporate a perceptual belief state accounted for dopamine signals^36,37^. Here, we confirm the involvement of dopaminergic RPEs in perceptual decision-making, but also show that this view is incomplete. In addition to signaling RPEs, dopamine release in the dorsal striatum also signals SPEs - violations of sensory rather than reward expectations. Critically, the TD-RL model with a perceptual belief state could not explain the opposite effects of reward size and stimulus probability on stimulus-evoked dopamine release. The model predicted stronger dopamine responses for both stimuli signaling larger rewards, and more probable and thus more frequently rewarded stimuli, because both increased expected reward value. It therefore lacked any mechanism for dopamine to increase when low-probability stimuli violated sensory expectations. This mismatch demonstrates that RPEs alone cannot account for the full pattern of stimulus-evoked dopamine modulation. Instead, a TD-RL model that, in addition to updating stimulus-choice values through RPEs, updates sensory priors through SPEs accounted for both similar behavioral biases and opposite stimulus-evoked dopamine signals. Notably, the stimulus probability manipulation also modulated dopamine signals at reward time with increased dopamine responses to rewards following low-probability stimuli. RPE signaling was sufficient to explain this modulation, as rare, and therefore infrequently rewarded stimuli induced lower reward expectations, leading to larger RPEs. Whether reward-evoked dopamine signal additionally reflects sensory surprise associated with the sensory features of reward cannot be determined from the present data and remains an important question for future work.

Previous studies have suggested that dopamine neurons send different types of prediction errors to different striatal regions, enabling distinct forms of learning across specialized neural circuits^40,55,56,68–70^. We here show that this spatial division does not universally hold: We observed both SPEs and RPEs in the same region of the dorsal striatum. This raises a key question: How can the striatum learn distinct functions for perceptual and reward learning from spatially overlapping dopamine signals, as proposed by our computational model? Recent theoretical models offer a potential solution^71,72^. In particular, if striatofugal projections undergo similar systematic long-term plasticity as corticostriatal projections, the principle of feedback alignment enables a demixing of multiple prediction errors downstream from the striatum, despite spatially diffuse dopamine signaling. Previously developed to explain the learning of multidimensional continuous actions^72^, and learning of multiple reward values according to different reward dimensions^71^, this principle may generalize to explain learning from overlapping SPEs and RPEs. Whether this overlap in RPEs and SPEs arises at the level of dopamine release or is already present in the activity of dopamine neurons remains an important question for future work.

Our results complement recent observations that pre-stimulus dopamine in the tail of the striatum reflects sensory expectations that bias immediate perception^46^. Those pre-stimulus signals represent predictions (priors) about the occurrence of sensory stimuli. In contrast, our findings identify the error term required to update those predictions. Dopaminergic SPEs may provide one of the mechanisms through which sensory predictions can be recalibrated. By revealing a causal and computational link between dorsal striatal dopamine release and perceptual learning, our results extend the role of dopamine from modulating perception in the moment to supporting the longer-term updating of sensory beliefs.

SPEs are ubiquitous in sensory cortices^24–28^, and predictive processing theories have predominantly focused on how cortical circuits perform inference and learning^23,29,30^. Our results point to an expanded framework for perceptual learning in which prediction error-driven updating is distributed across cortical and subcortical circuits and shaped by neuromodulators. This is supported by two lines of evidence. First, large-scale electrophysiology recordings show that prior information during perceptual decision-making is present throughout the mouse brain^50,52^. Second, complementary work has shown that noradrenergic neurons in the locus coeruleus broadcast unsigned SPEs across the cortex^73^, suggesting a broader role of neuromodulators in perceptual learning. Future work integrating these neuromodulatory and subcortical mechanisms with cortical predictive processing frameworks will be crucial for establishing a unified theory of how the brain generates, updates, and maintains perceptual expectations.

We found that SPE signaling was specific to dopamine and absent in striatal acetylcholine. This is surprising, as striatal dopamine and acetylcholine signals are typically tightly coupled^64,74^. Our data reveal two key dissociations between these signals. First, dopamine responded bilaterally to sensory input, whereas acetylcholine responses were lateralized to contralateral visual stimuli. Second, only dopamine was modulated by sensory surprise, while acetylcholine was not. This dissociation complements prior observations that putative cholinergic interneurons in the striatum do not encode reward expectations^75^. This suggests that both RPE and SPE signaling are specific to dopamine, whereas acetylcholine provides more veridical, context-independent visual information. The functional significance of this distinction remains to be determined. Importantly, while previous work has shown that acetylcholine can elicit local dopamine release^62^, our results suggest that such mechanisms do not account for dopaminergic SPEs.

Our study has a number of limitations. First, our fiber photometry recordings measure bulk dopamine release at coarse spatial resolution. Our recordings therefore cannot reveal potentially more fine-grained spatial organization in dopaminergic SPEs and RPEs. For instance, while our dopaminergic SPEs is best understood as unsigned prediction errors, signaling absolute, rather than stimulus-specific surprise, it is possible that dopamine encodes signed prediction errors at finer spatial scales. Notably, such signed errors are not strictly required for stimulus-specific learning. Instead, the content of prediction errors could be encoded in cortical inputs to striatal neurons, with dopamine strengthening synaptic plasticity only at co-active corticostriatal synapses^76^. This mechanism provides a plausible explanation for why broad, non-specific optogenetic stimulation of dopamine release at stimulus onset produced stimulus-dependent choice biases in our experiment. Second, our recordings were restricted to the DLS and to a visual decision-making task. Whether SPEs and RPEs are more or less segregated in other striatal regions, and whether they are sensory-modality specific, remain open questions.

In summary, our results show that dopamine in the same dorsal striatal circuit conveys multiple teaching signals, supporting distinct forms of learning. Our finding that dopamine encodes SPEs highlights a departure from the traditional reward-centric view, pointing to a broader role of dopamine in shaping perceptual inference and decision-making.

## Methods

### Animals

The data presented in this paper was collected from 29 male wild-type C57/BL6J and DAT-Cre mice, with their age ranging between 10 to 30 weeks. The first 13 animals were wild-type mice trained on the visual decision-making task. Eight of these thirteen mice expressed dopamine and acetylcholine sensors in opposite dorsolateral striata (DLS). Two and three animals only expressed dopamine or acetylcholine sensors in one hemisphere, respectively. Of the 13 animals, 8 performed the stimulus probability manipulation, 2 performed the reward size manipulation, and 3 performed both manipulations. The visual classical conditioning task was performed by 6 wild-type mice with bilateral DLS dopamine recordings. The optogenetic dopamine stimulation experiment involved 10 DAT-Cre mice, expressing ChrimsonR. Mice were kept on a 12 h dark/light cycle, with an ambient temperature of 20–24° Celsius, and 40% humidity. All experiments were conducted according to the UK Animals Scientific Procedures Act (1986) under appropriate project and personal licenses.

### Surgery

Animals were anaesthetized with isoflurane and were kept on a feedback-controlled heating pad (Stoelting 53810). Hair overlying the skull was shaved and the skin and muscles over the central part of the skull were removed. The skull was thoroughly washed with sterile saline. A head plate was attached to the bone posterior to bregma using dental cement (Super-Bond C&B). After the head plate fixation, we made craniotomies over the target areas and injected 450nl of AAV9-hsyn-GRAB_DA2m (for recording dopamine) or 450nl of AAV9-hSyn-GRAB-ACh3.0 (for recording acetylcholine) into the right and/or left DLS (AP: +0.5mm from bregma; ML: +/-2.5mm from midline; DV: 2.8mm from dura). For the optogenetic experiments, we injected 150nl of AAV5-Syn-FLEX-rc[ChrimsonR-tdTomato] into the right and left SNc (AP: -3mm; ML: +/-1.5mm; DV: -4.3mm) of DAT-Cre mice. This was followed by implantation of optic fibers over the DLS (core = 200 um, Neurophotometrics Ltd; tip 0.3 mm above the injection site for recording experiments). The fibers were secured to the head plate and skull using dental cement. Mice recovered for at least 5 days following the surgery. We waited for an additional 3 weeks for GRAB sensor expression and 4-5 weeks for opsin expression in mice with ChrimsonR injections, before recording and stimulation experiments.

### Materials and apparatus

Mice were trained on a standardized behavioral rig, consisting of an LCD screen (9.7” diagonal), a custom 3D-printed mouse holder, and a head bar fixation clamp to hold a mouse such that its forepaws rested on a steering wheel^49,77^. Silicone tubing controlled by a pinch valve was used to deliver water rewards to the mouse. The general structure of the rig was constructed from Thorlabs parts and was placed inside an acoustical cabinet. The experiments were controlled by freely available custom-made software^78^, written in MATLAB (Mathworks). Data analyses were performed with custom-made software written in Matlab 2020b and R (version 4.3.1).

### Visual decision-making task

Behavioral training in the visual decision-making task started at least 5 days after the surgery. Animals were handled and acclimatized to head fixation for at least 3 days, and then trained in a 2-alternative forced choice visual detection task^77^. After mice kept the wheel still for at least 0.7 to 0.8 s, a sinusoidal grating stimulus of varying contrast appeared on either the left or right side of the screen (±35° azimuth, 0° altitude). Grating stimuli had a fixed vertical orientation, were windowed by a Gaussian envelope (3.5° s.d.), and had a spatial frequency of 0.19 cycles/° with a random spatial phase. Concomitant to the appearance of the visual stimulus, a brief tone was played to indicate that the trial had started (0.1 s, 5 kHz). Mice were able to move the grating stimulus on the monitor by turning a wheel located beneath their forepaws. If mice correctly moved the stimulus 35° to the center of the screen, they immediately received a water reward (2–3 μL). Conversely, if mice incorrectly moved the stimulus 35° towards the periphery or failed to reach either threshold within 60 s a noise burst was played for 0.5 s and they received a timeout of 2 s. The inter-trial interval was randomly sampled from a uniform distribution between 0.5 and 1 s (1 and 3 s in subsequent recording experiments). In the initial days of training, only 100% contrast stimuli were presented. Stimuli with lower contrasts were gradually introduced after mice exhibited sufficiently accurate performance on 100% contrast trials (>70% correct). During this training period, incorrect responses on easy trials (contrast 50%) were followed by “repeat” trials, in which the previous stimulus location was repeated. The full task included six contrast levels (100, 50, 25, 12.5, 6.25 and 0% contrast). Once mice reached stable behavior on the full task, repeat trials were switched off, and mice proceeded to the main experiment.

In the main experiment (**Fig. 1**, **2**, and **4**), we investigated whether mice adapt their visual decisions to asymmetric reward sizes and biased stimulus probabilities. In the reward size manipulation, mice received a larger (3 uL) water reward for correctly reporting one stimulus size, and received a smaller reward (1.5 uL) for correctly reporting the other side. For 3 mice the larger reward size was switched several times per session, in blocks of 70 to 90 trials. For 2 mice the larger reward size was switched across recording days. No reward was given for incorrect choices and stimuli appeared with equal probability on both sides. In the stimulus probability manipulation, mice received a symmetric water reward (2 uL) for correctly reporting either stimulus location, but stimuli were more likely to appear on one side (80 vs 20% probability). The more likely stimulus side was switched several times per session, in blocks of 10 to 100 trials.

### Visual classical conditioning task

In the visual classical conditioning task we sought to test whether sensory prediction errors (SPEs) would occur in the absence of instructed actions. We trained naive mice to associate visual stimuli with water reward. We started by presenting a large (s.d. = 20) drifting grating (drift speed = 1.5 cycles/s) for one second at the center of the screen. Concomitant with the offset of the grating, mice received a water reward (3 uL). Mice did not have access to the response wheel, and reward was not contingent on the mice’s behavior. The subsequent visual stimulus was presented after an inter-trial interval of 6 to 60 seconds, randomly sampled from a truncated exponential distribution. Over several days of training, we progressively decreased the size and drift speed of the gratings, until gratings had a Gaussian envelope of 3.5° s.d. and were static. Furthermore, we gradually moved gratings into the periphery (±35° azimuth). During training, gratings appeared on the left and right side with equal probabilities. We subsequently manipulated the probability of stimuli appearing on the left or right side of the screen across recording days (80 vs 20% probability), while keeping all other parameters constant. We monitored uninstructed behavior, such as anticipatory and consummatory licking, and changes in pupil size through video recordings, as described below.

### Video monitoring

The left eye was monitored with a camera (Flir CM3-U3-13Y3M-CS) fitted with a zoom lens (Thorlabs MVL50M23) recording at 20 Hz. Front body movements were monitored with another camera (same model but different lens, Thorlabs MVL16M23) also recording at 20 Hz. Mice were illuminated with infrared light (850nm, BW BWIR48) for the recording of eye and front body movements.

### Dopamine and acetylcholine recordings

To measure dopamine or acetylcholine release, we employed chronic fiber photometry^79,80^. We used multiple excitation wavelengths (470 and 415 nm), delivered on alternating frames (sampling rate of 40 Hz), serving as target and isosbestic control wavelengths, respectively. We collected emitted fluorescence through the same implanted fibers. To remove movement and photobleaching artifacts, we subtracted the isosbestic control from the target signal. In particular, for each session, we computed a least-squares linear fit of the isosbestic to the target signal. We subtracted the fitted isosbestic from the target signal and normalized by the fitted isosbestic signal to compute ΔF/F:

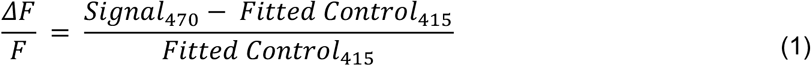

The resulting signal was further high-pass filtered by subtracting a moving average (25 s averaging window) and z-scored.

### Dopamine optogenetic stimulation

We trained mice expressing the excitatory opsin ChrimsonR in the DLS dopamine terminals on the visual decision-making task. Upon reaching a stable accuracy above 65%, we began the optogenetic experiment. On 40% of randomly interleaved trials, laser pulses (25ms on/25ms off, 635nm, 7mW measured at the tip of the patch cord) were delivered through the optic fibers over a period of 250 ms at the onset of visual stimuli. In order to collect more trails per visual contrast, we reduced the set of contrasts to 0, 25 and 50%. The inter-trial-interval was randomly sampled from a uniform distribution between 2 and 3 seconds.

### Histology and fiber track quantifications

Histology was performed after the experiments to confirm successful fiber positioning. Animals were deeply anaesthetized and perfused using 4% paraformaldehyde (PFA) and then decapitated. The brains were extracted, left in 4% PFA for 24h to post-fix in a refrigerator and then embedded in blocks of 1.5% agarose gel before collecting slices at 70 um thickness using a vibratome (Leica VT1000 S). Slices were then stained with DAPI for 15 min (1:1000 solution), mounted onto glass, coverslipped, and imaged using an epifluorescence microscope (Zeiss AxioZoom).

### Quantification and statistical analysis

#### Behavioral analyses

We excluded experimental sessions in which behavioral performance was poor using the following criteria. We fit a psychometric curve to each session’s data, using a maximum likelihood procedure:

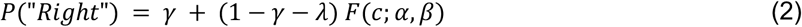

where P(“Right”) describes the mouse’s probability to give a rightward response, F is the logistic function, *c* is the stimulus contrast, *γ* and *λ* denote the right and left lapse rates, *α* is the bias and *β* is the contrast threshold. We excluded sessions in which the absolute bias was larger than 0.16, or either left or right lapse rates exceeded 0.3. We further excluded sessions in which the choice accuracy on easy 100% contrast trials was lower than 80%. This led to the exclusion of 66 out of 273 sessions (24%). Finally, we excluded trials in which the response time was longer than 12 s, thereby excluding 518 out of 87,625 trials (0.6%).

To assess how perceptual decisions adapted to the reward size and stimulus probability blocks, we plotted the proportion of rightward responses, P(“Right”), as a function of stimulus contrast, conditioning this proportion on the current reward size block (left or right large reward) or stimulus probability block (left or right likely). We summarized choice bias as the difference in P(“Right”) between right and left blocks. A positive ΔP(“Right”) indicated a tendency to choose the large-reward or high-probability side. To statistically assess biases, we averaged ΔP(“Right”) across stimulus contrasts, and compared the average bias to zero, using one-sample two-sided t-tests. To compare the biases across manipulation, we used a two-sample two-sided t-test. The critical alpha thresholds were set at 0.05.

#### Pupil analyses

We used DeepLabCut^81^ to track several points on the mice’s left pupil throughout each task trial. We selected 4 points in the top, bottom, left and right portions of each mouse’s pupil and recorded the x and y coordinates of each point over time. We defined the pupil diameter as the mean of the Euclidean distances between the top and bottom and left and right points. The time series of pupil diameter was z-scored per session. To evaluate the modulation of pupil size by stimuli of different probability, or reward size, we first time-locked pupil responses to stimulus onsets and calculated the average pupil size between 1 and 3 s after stimulus onset for each trial. We then divided the pupil size on high-probability or large-reward trials by that on low-probability or small-reward trials to obtain a modulation index. We performed one-sample t-tests, comparing the log-transformed modulation indices against zero, as is appropriate for response ratios. The critical alpha threshold was 0.05. We used the same procedure for the visual decision-making and classical conditioning tasks.

#### Lick analysis

We used FaceMap^82^ to track the mouth and lower lip key points on videos of the mice’s front body throughout each task trial. We recorded the x and y coordinate of each key point and defined a mouth-lip (Euclidean) distance measure that served as a proxy of licking activity. To evaluate differences in anticipatory, pre-reward licking for low vs. high probability stimuli in the classical conditioning task, we calculated the trial-by-trial average licking activity during stimulus presentation (0 to 1 s) before reward delivery. We then divided the licking activity for high-probability stimuli by that for low-probability stimuli to obtain a modulation index. We statistically tested for differences using a one-sample t-test, comparing the log-transformed modulation index against zero.

#### Linear encoding model of dopamine and acetylcholine release

To assess which task events contributed to variations in dopamine and acetylcholine release in the DLS, we used linear encoding models (linear deconvolution). This model regresses binary variables indicating the time period around events onto full trial fluorescent signals. The design matrix (*X*) has a column per time point around the events of interest (regressors) and a row per time point of the fluorescent signal, which was concatenated for all trials across all sessions for a given animal. The concatenated fluorescent signal was used in the regression to form the dependent variable (*y*). Model fitting amounts to finding the optimal scaling (*β*′) of the regressors in *X* which produces a prediction of the concatenated fluorescent signal (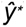) that minimizes the mean squared error 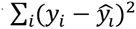. The least squares solution is given by:

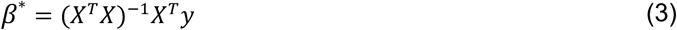

the prediction of the fluorescent signal is given by

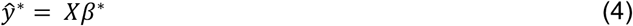

The resulting elements in *β*^∗^ compose the deconvolved signal for each event. The benefits of using linear deconvolution over an event-aligned average is that it isolates the effect of each event on the fluorescent signal, removing the influence of other events occurring shortly before or after. This isolation is achieved as long as events are sufficiently decorrelated, e.g., due to temporal jitter. The full model comprised the following events, each modeled from 0.5 to 2 s, relative to event onset: Stimulus events per visual contrast on each side, expressed relative to stimulus onset (11 events), separate choice events for left and right wheel turns, per visual contrast, expressed relative to choice onset (11 events), reward events per visual contrast, expressed relative to reward delivery (11 events). We discarded unrewarded trials prior to model fitting. We also fit an alternative model with binary choice events (separate left and right wheel turns, independent of visual contrast), but this had little impact on model fits. We subsequently fit reduced models, by dropping one of the three events groups (stimulus, choice or reward events). To assess model fits, we computed each model’s R^2^, using leave-one-session-out cross-validation. R^2^ was computed on the held-out test data as

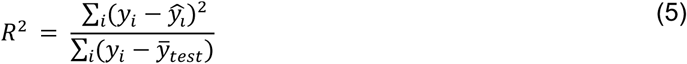

where 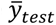 is the mean of all *y_i_* in the test set. Finally, we computed the reduction in R^2^ for each reduced model relative to the full model as

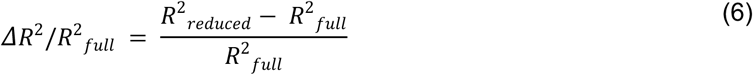

#### Dopamine and acetylcholine event-related analyses

We aligned the z-scored ΔF/F fluorescent signals to stimulus or reward onset times. When analyzing stimulus-evoked responses, we baselined the signal to the pre-stimulus period of the current trial (-0.5 to 0 s relative to stimulus onset). For dopamine recordings in the decision-making task, trial-by-trial stimulus-evoked responses were calculated as the mean response 0.2 to 0.4 s after stimulus onset (excluding time points after reward). For acetylcholine, we separately analyzed early peak responses (0.1 to 0.3 s) and subsequent troughs (0.3 to 0.5 s). When analyzing reward-evoked responses, we baselined the signal to -0.1 to 0 s relative to reward. For dopamine recordings in the decision-making task, trial-by-trial reward-evoked responses were calculated as the mean response 0 to 1 s after reward. For acetylcholine, we separately analyzed early peak responses (0 to 0.2 s) and subsequent troughs (0.2 to 1 s). Unless otherwise noted, we analyzed rewarded trials only, to ensure that mice were engaged with the decision-making task. In the visual classical conditioning experiment, average stimulus- and reward-evoked dopamine responses were calculated from 0 to 1 s (stimulus presentation) and 1 to 2 s relative to stimulus onset.

To evaluate the modulation of dopamine and acetylcholine by stimuli of different probability, or reward size, we divided the mean stimulus responses on high-probability or large-reward trials by those on low-probability or small-reward trials to obtain modulation indices. We performed one-sample t-tests, comparing the log-transformed modulation indices against zero, as is appropriate for response ratios. To assess differences between manipulations, we performed a two-sample t-test comparing log-transformed modulation indices. The critical alpha thresholds were 0.05. We performed analogous analyses on reward responses, but focused on trials in which no stimulus was presented, but the mouse reported the low/high stimulus probability or reward size side. This ensured that reward responses were not confounded by overlapping stimulus responses. However, we observed similar modulations for reward responses on trials with visual stimuli (**Fig. S2h,r** and **S4o**).

To further probe whether dopamine and acetylcholine were primarily modulated by stimulus processing or choice execution, we performed control analyses in which we binned trials into sextiles according to response times, separately for each contrast level. This ensured that bottom-up stimulus contrast was matched across bins, while bins systematically varied in their response times (**Fig. S2e,o** and **S3e**). If neuromodulator responses would be time-locked to choice execution, one would expect to observe progressively later responses with increasing response times. To test this prediction, we plotted fluorescent responses, split by response time bin, and quantified peak response magnitudes and latencies (**Fig. S2f-g,p-q** and **S3f-i**). To test for systematic linear trends across mice we fit linear models to peak response magnitudes and latencies as a function of bin and compared the slope parameters against zero using one-sample t-tests.

#### Dopamine optogenetic stimulation analysis

We excluded sessions in which choice accuracy on 50% contrast trials was below 65%, removing 23 of 113 sessions (20%). Trials with response times longer than 12 s were also excluded. To assess how optogenetic stimulation of dopamine release at stimulus onset influenced subsequent perceptual decisions, we analyzed specific trial pairs. The preceding trial met three criteria: it was rewarded, had a stimulus contrast of 0% or 50%, and was either optogenetically stimulated or not. The subsequent (current) trial had no stimulation and could have any contrast. This design allowed us to measure perceptual decisions in the absence of stimulation, depending on whether the previous stimulus had been paired with optogenetically driven dopamine release. Trials with previous stimulation at zero contrast served as a control condition, since such trials were not expected to update priors in a directional manner. Decision bias was quantified as the difference in the proportion of rightward choices between trials preceded by right- versus left-stimulus trials, ΔP(“Right”). A positive ΔP(“Right”) indicated a bias toward choosing the side of the previous stimulus. We compared ΔP(“Right”) between previously stimulated and non-stimulated trials using a paired t-test. To assess the temporal decay of stimulation-induced decision bias, we repeated the analysis while conditioning on stimulation occurring 2 to 41 trials earlier. The resulting biases were averaged in windows of 10 trials. For each window, we used one-sample t-tests with Bonferroni correction to assess statistical significance.

#### Ideal observer models

We developed a normative agent that simultaneously engages in Bayesian perceptual inference and reinforcement learning. We sought to test whether this agent would jointly account for our mice’s similar behavioral biases but opposite dopamine modulations across the reward size and stimulus probability manipulations. To this end, we adopted and extended a previously proposed Reinforcement Learning model based on a partially observable Markov decision process (POMD)^37,66^. We first describe this reinforcement learning model, hereafter referred to as the *RL agent*, and then outline a perceptual inference (*PI*) agent, and finally describe the combination of the two processes (*RL + PI*).

##### RL agent

In our visual decision-making task, the state of the current trial (left or right) is uncertain and therefore only partially observable due to the presence of low contrast stimuli and sensory noise. We assume that the agent forms an internal estimate 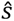 of the true signed stimulus contrast *s*, which is normally distributed with constant variance around the true stimulus contrast: 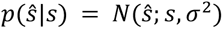. The sensory evidence for the stimulus being presented at the right side of the monitor is given by:

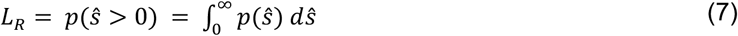

where *L* is the sensory likelihood. We assume that the agent maintains a neutral (uniform) expectation about which side the stimulus is presented. Therefore, the agent’s posterior belief that the stimulus was presented on the right side of the monitor is given by:

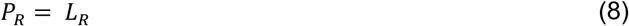

We note that this equality is specific to an agent with neutral prior expectations, and will not hold for agents with history-dependent priors (see below). The agent’s belief that the stimulus was presented on the left side is given by:

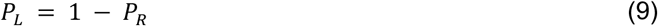

The agent combines this belief state [*P_L_*, *R_R_*] with stored values for making left and right choices in left and right perceptual states, given by *q_choice,state_* in order to compute expected values for left and right choices:

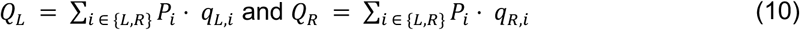

where *Q_L_* and *Q_R_* denote the expected value for left and right choices, respectively.

The agent then uses these expected values together with a softmax decision to compute a probability of making a rightward choice, P(“Right”):

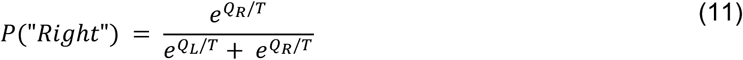

where *T* denotes the softmax temperature, introducing decision noise. The agent then decides for one of the two choice options using a biased coin flip, based on P(“Right”).

Following each choice, the agent observed the corresponding outcome *r*, which was 1 for a correct small reward trial and 0 for incorrect trials. To model subjective valuation, large rewards were multiplied by a subjective reward scaling factor 1 ≤ *S_R_* ≤ 2 , capturing potential diminishing marginal utility. The agent then computes a reward prediction error (RPE) 𝛿 by comparing the outcome to the expected value of the chosen option *QC*:

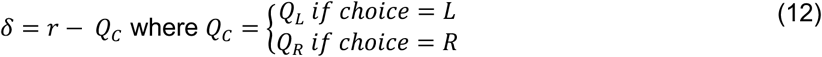

Given this prediction error, the agent updates the values associated with the chosen option *q_C,L_* and *q_C,R_* by weighing the prediction error with a learning rate *α_RL_* and the belief of having occupied the particular state:

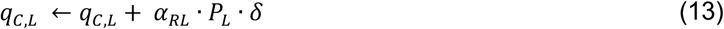

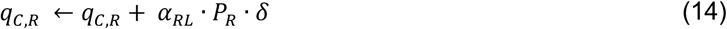

The RL agent had four free parameters, consisting of sensory noise *σ*^2^, decision noise *T*, reinforcement learning rate *α_RL_*, and subjective reward scaling factor *S_R_*.

##### PI agent

We next developed an agent that performs perceptual inference and the absence of reward-based learning, serving as a control model. This perceptual inference (PI) agent computes a sensory likelihood, *L_R_*, analogously to the RL agent (equation 7). However, the agent maintains a sensory prior expectation, defined as an estimate of how frequently a stimulus appears. Specifically, the agent forms the sensory prior by heuristically tracking stimulus base rates^50,52^. This is achieved by computing a trial-by-trial SPE ɣt, as the difference between received and expected sensory input:

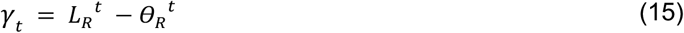

and updating the sensory prior with this predictions error, weighted by a perceptual learning rate *α_Perception_*. Following previous studies^50,83^, we also incorporated a common finding in the literature known as conservatism (i.e., observers are biased towards a uniform prior), by averaging the prior with a uniform prior of 0.5:

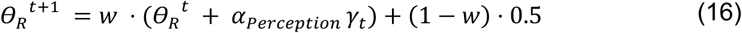

The prior expectation for stimuli on the left is given by *θ_L_* = 1 − *θ_R_*. To infer the stimulus side, the agent combines the noisy sensory measurement *L_R_^t^* (likelihood) with the sensory prior *θ_R_^t^* according to Bayes rule, resulting in a posterior belief state:

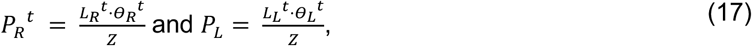

Where Z denotes a normalization constant, chosen such that *P_R_^t^* and *P_L_^t^* sum to 1. The belief about the stimulus side is therefore not only dependent on the sensory measurement, but also on the agent’s prior expectation. To remain consistent with the RL agent, and to introduce post-inference decision noise, the probability of making a rightward choice, P(“Right”), was then computed by combining the posterior belief state with a softmax decision rule:

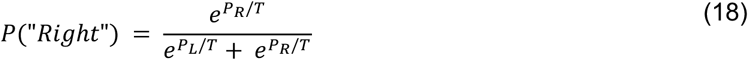

where *T* denotes the softmax temperature, introducing decision noise. The agent then decides for one of the two choice options using a biased coin flip, based on P(“Right”), and thereby tends to choose the option that is more likely. The PI agent had four free parameters, consisting of sensory noise *σ^2^*, decision noise *T*, perceptual learning rate *α_Perception_*, and conservatism weight *w*.

##### PI + RL agent

The PI + RL agent simultaneously engaged in the perceptual inference and reinforcement learning processes described above. That is, perceptual belief states were computed using Bayesian inference (equations 7 and 17), and sensory priors were updated through sensory prediction errors (equations 15 and 16). The resulting posterior belief states were used for value-based decision-making (equations 10 and 11) and reward-based learning (equations 12-14). The PI + RL agent had 6 free parameters, consisting of sensory noise *σ^2^*, decision noise *T*, perceptual learning rate *α_Perception_*, conservatism weight *w*, RL learning rate *α_RL_* and subjective reward scaling factor *S_R_*.

##### Fitting procedure and model comparison

We fit all three models to the joint behavioral choice data of the stimulus probability and reward size manipulations pooled across mice. The likelihood of the empirically observed choices, *c_t_* ∈ {*L, R*}, under a given model was

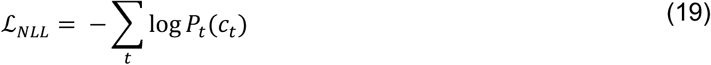

Where the model’s probability for generating the same choice as the mouse was given by

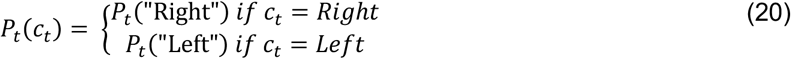

The model’s probability for a rightward choice on trial t was calculated using equations 11 or 18, and *P_t_* ("Left") = 1 − *P_t_*("Right"). In order to increase the stability of the parameter search, for each function evaluation (equation 19), we simulated the agent 100 times, and averaged the model’s trial-by-trial choice probabilities 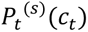 across all simulations s:

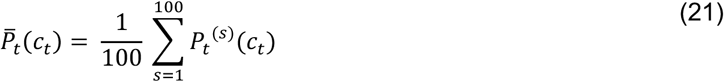

We used these average choice probabilities (equation 21) to compute negative log-likelihoods^84^:

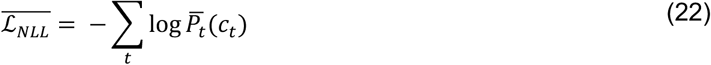

We minimized the negative log-likelihood (equation 22) using a Bayesian adaptive direct search method^85^ . This method alternates between a series of fast, local Bayesian optimization steps and a systematic, slower exploration of a mesh grid. We constrained the parameter search space as follows. The sensory noise parameter was constrained to the interval [0.05, 0.5], decision noise to [0.01,1], perceptual and RL learning rates to [0.01, 1], conservatism weight to [0,1], and the subjective reward scaling factor to [1,2]. We repeated the parameter search process from multiple different starting points to confirm converging solutions. Perception and RL learning rates, and conservatism weight were initialized with 0.2 or 0.8, and the subjective reward scaling factor was initialized with 1.2 or 1.8. Sensory and decision noise were initialized with 0.1 and 0.3 respectively. The different starting parameter combinations yielded 16 starting points for the PI + RL agent, and 4 for RL and PI agents. We compared the best-fitting models using likelihood ratio tests, as well as the Bayesian information criterion (BIC). The best fitting parameters for the PI + RL agent are shown in **Fig. S6a**.

##### Model predictions for dopamine responses

We used the best-fitting behavioral models to derive predictions for empirical dopamine responses, based on the model’s latent variables. First, we assumed that dopamine responses of the RL agent would encode a temporal difference RPE that evolves throughout the trial (see **Fig. 6b**). At the time of stimulus processing, the model’s dopamine encoded the expected reward, before committing to a choice, given by

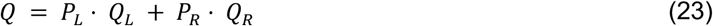

At the time of outcome, the models’ dopamine encoded the RPE 𝛿 (equation 12). Second, we assumed that the dopamine responses of the PI agent would encode an unsigned SPE |𝛾| (see equation 15) at the time of stimulus processing, i.e., the absolute deviation of sensory measurements from prior expectations. Finally, we assumed that dopamine responses of the RL + PI agent would reflect a weighted linear combination of temporal difference RPEs and SPEs (**Fig. 6b**).

To evaluate the congruence between empirical and model dopamine predictions, we fit the model-derived dopamine predictions to the empirical dopamine stimulus responses across the reward size and stimulus probability manipulations (averaged between 0.2 and 0.4 post-stimulus, correct trials only). We allowed for different intercepts and general scaling coefficients across the two manipulations, as they were recorded in partially distinct groups of animals. However, we assumed that there is a shared linear mixing coefficient, determining the overlap of TD-RPEs (Q at the time of stimulus presentation; equation 23) and SPEs:

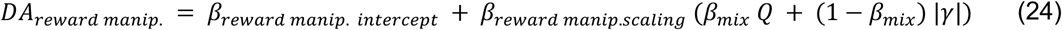

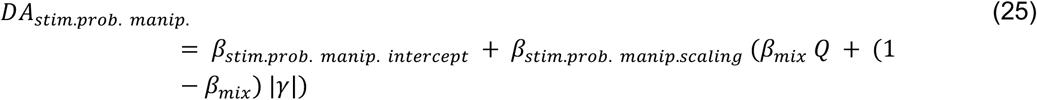

Predictions for reward responses were generated using the same intercept, scaling and mixture coefficients. The best fitting mixture coefficient *β_mix_* for the PI + RL agent was 0.55.

## Acknowledgements

This work was supported by grants from the Wellcome Trust (213465/Z/18/Z) and ERC (funded by UKRI, EP/X026655/1) to A.L. M.F. is supported by a HFSP long-term postdoctoral fellowship (LT0045/2022). R.B. is supported by grants from BBSRC (BB/S006338/1) and MRC (MC_UU_00003/1).

## Author Contributions

M.F. and A.L. conceived the study. M.F., C.T. and A.L. designed the experiments. M.F., C.T., Y.S., O.D., R.P., and A.M. performed the experiments and acquired the data. M.F. performed the formal analysis, with inputs from R.B. and A.L. M.F. and A.L. interpreted the results, with contributions from R.B. M.F. and A.L. wrote the manuscript, with contributions from R.B.

## Additional information

**Correspondence and requests for materials** should be addressed to Matthias Fritsche or Armin Lak.

---

**Fig. S1.**
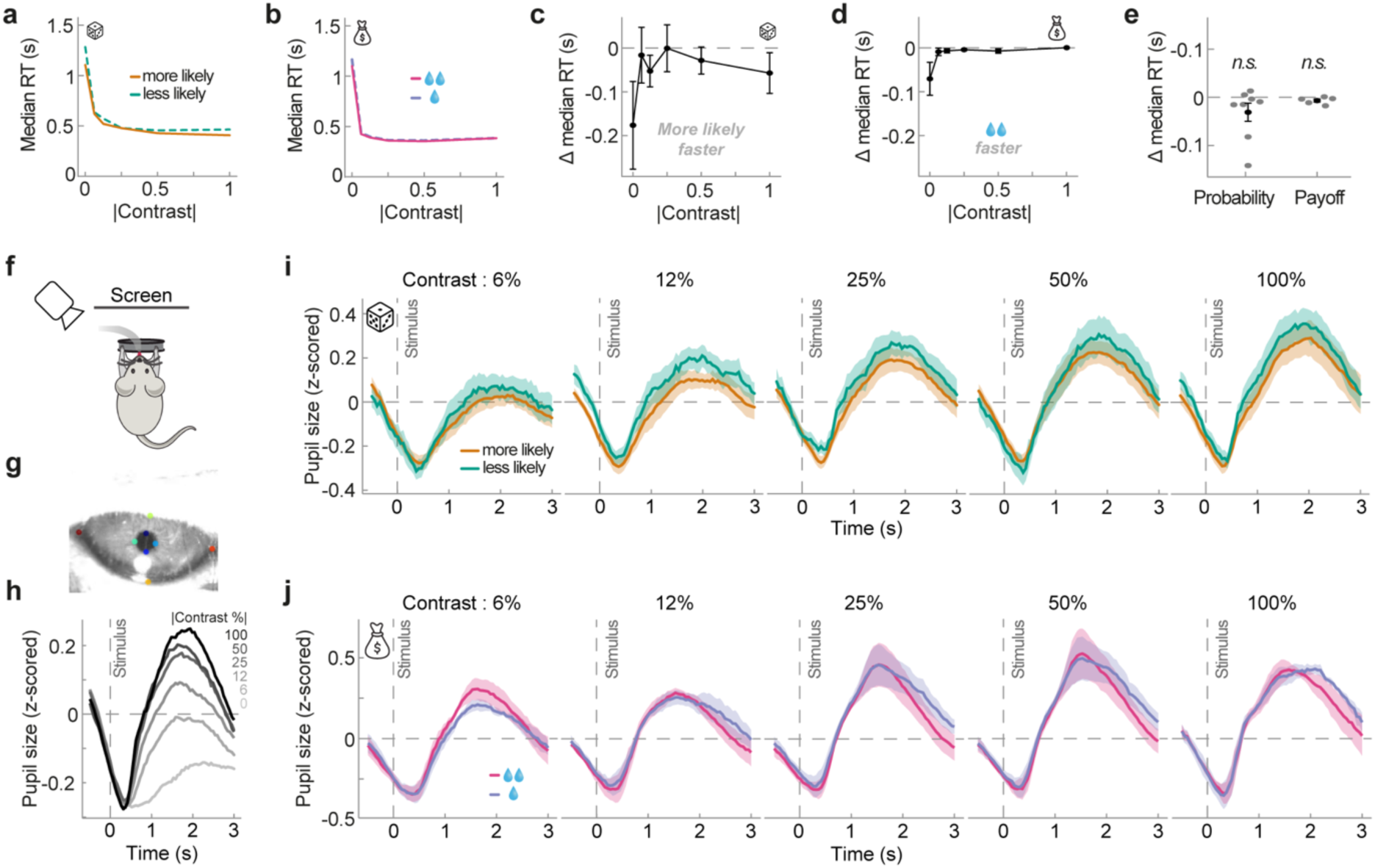
Response times and pupil responses in the visual decision-making task. **a,** Median response times as a function of absolute visual stimulus contrast (x-axis) and stimulus probability (color). Lines show averages across animals. Correct trials only. **b,** Same as in panel a, but for reward size manipulation. **c,** Difference in median response times between high and low probability stimuli, per absolute stimulus contrast. Negative y values indicate faster responses to more likely stimuli. Data points show means across animals, and error bars denote SEMs. **d,** Same as in panel c, but for reward size manipulation. Negative y-values denote faster responses to stimuli signaling larger reward. **e,** Response time differences in probability and reward manipulations, averaged across stimulus contrasts. There was no significant effect of either stimulus probability or reward size. Grey dots denote individual animals, black dots show group average. Error bars denote SEMs. **f,** Schematic of the video recording setup. **g,** Example video frame of the eye recording with DeepLabCut markers (colored dots). **h,** Group-average pupil responses to visual stimuli with different absolute contrasts. Post-stimulus pupil dilation increases for higher contrast stimuli. **i,** Group-average pupil responses for more (orange) and less likely stimuli (green). Different panels show different stimulus contrasts. Shaded regions denote SEMs. **j,** Same as in panel i, for more (pink) and less rewarding stimuli (blue). Pupil responses revealed a selective effect of probability: surprising stimuli presented on the low-probability side evoked larger post-stimulus pupil dilations than expected stimuli (t(8) = 3.54, p = 0.009; panel i). In contrast, reward asymmetry did not significantly affect pupil size (t(4) = 0.50, p = 0.64; panel j). See also Fig. 1.

**Fig. S2.**
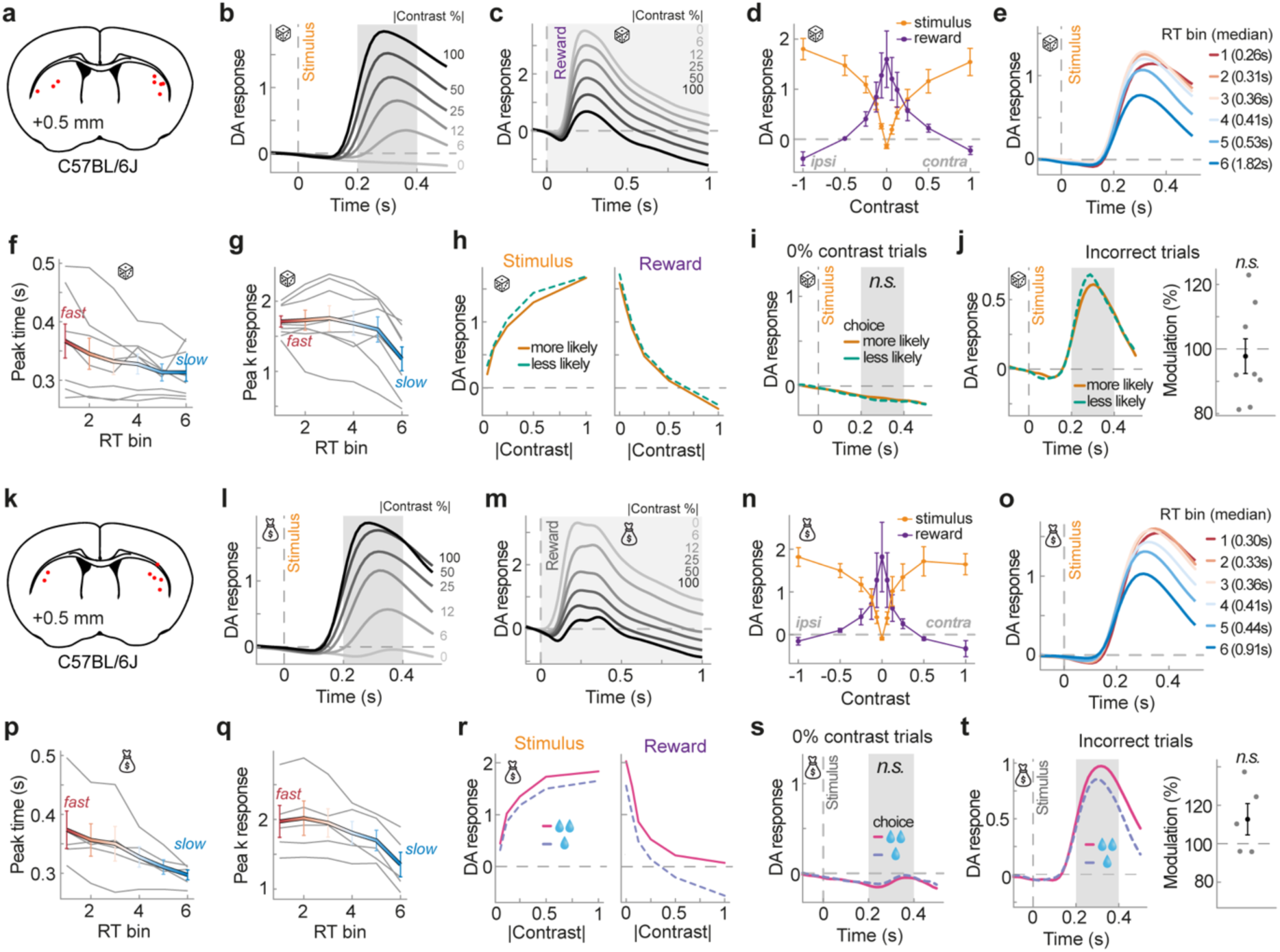
Dopamine recordings in the stimulus probability and reward size manipulation. **a,** The locations of DLS optic fiber implants for dopamine recordings in mice performing the visual decision task with stimulus probability manipulation, estimated from post-mortem histological examinations overlaid on a brain slice schematic. **b,** Group-average dopamine response aligned to stimulus onset (gray dashed line) in the stimulus probability manipulation, split by absolute stimulus contrast (gray to black; correct trials only). Gray shaded area indicates the stimulus time period over which we averaged stimulus responses (d,h,i-j; excluding time points after reward delivery). **d,** Group-average stimulus-evoked (yellow) and reward-evoked (purple) dopamine responses in stimulus probability manipulation as a function of signed stimulus contrast relative to the recorded hemisphere (rewarded trials only; averaged over gray shaded areas in b). Positive x values denote stimuli contralateral to the recorded hemisphere. Dopamine responds similarly for ipsi- and contra-lateral stimuli. Error bars denote SEMs. **e,** Group-average stimulus-evoked dopamine response in stimulus probability manipulation, split into six response time bins (red to blue), while controlling for stimulus contrast. Bin 1 contains the 1/6 fastest responses of each contrast level. **f,** Peak time of dopamine responses as a function of response time bin. Peak time decreases with increasing response time (t(7) = -3.18, p = 0.015). Grey lines show individual mice. The thick colored line shows the group average. Error bars denote SEMs. **g,** Same as in panel f, but for peak height. Peak height decreases with increasing response times (t(7) = -3.81, p = 0.007). **h,** Stimulus- (left) and reward-evoked (right) dopamine responses for more (orange) or less likely stimuli (green), as a function of absolute stimulus contrast. Lines denote group averages. **i,** Group-average stimulus-locked dopamine response on 0% contrast trials, split by whether mice chose the more (orange) or less likely side (green). There was no significant effect of probability on these trials without visual stimuli. **j,** Stimulus-evoked dopamine responses on incorrect trials split by stimulus probability. There was a trend towards larger responses to low probability, surprising stimuli, which did not reach significance. This analysis is likely underpowered as incorrect, low- and high-probability stimuli only constitute 3% and 9% of all trials. **k-t,** same as panels a-j, but for reward size manipulation. Panel p: Dopamine peak time tends to decrease with increasing response time (t(4) = -2.69, p = 0.055). Panel q: Dopamine peak height decreases with increasing response time (t(4) = -4.68, p = 0.009). See also Fig. 2.

**Fig. S3.**
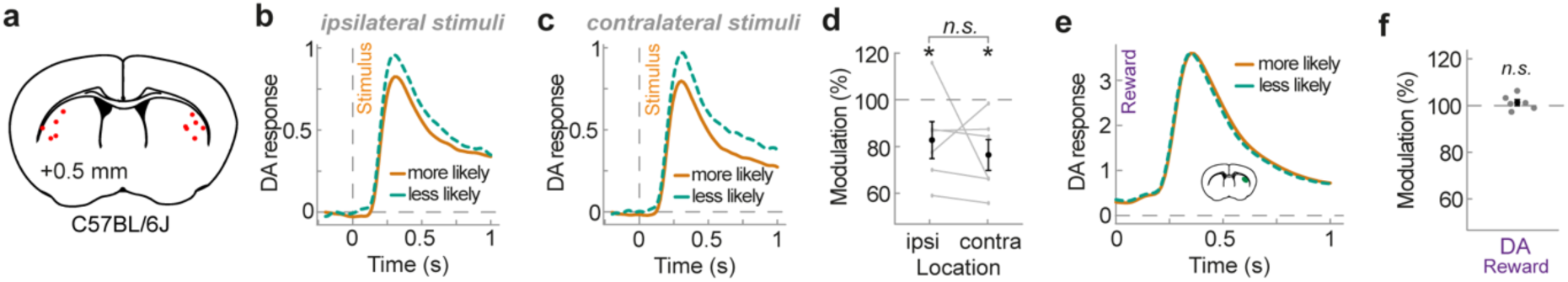
Dopamine recordings in the visual classical conditioning task. **a,** The locations of DLS optic fiber implants for mice performing the visual classical conditioning task, estimated from post-mortem histological examinations overlaid on a brain slice schematic. **b,** Group-average stimulus-evoked response for stimuli ipsilateral to the recorded hemisphere, split by stimulus probability (orange vs. green). **c,** Same as panel b, for contra-lateral stimuli. **d,** Modulation of responses by stimulus probability, separately for ipsi- and contra-lateral stimuli. Low probability, surprising stimuli evoked larger dopamine responses than high probability, expected stimuli, regardless of side (contra: t(5) = -3.29, p = 0.011; ipsi: t(5) = -2.26, p = 0.037, one-sided one-sample t-tests; contra vs. ipsi: t(5) = 0.72, p = 0.50, two-sided paired t-test). **e,** Reward-evoked dopamine responses, split by stimulus probability. **f,** Reward-evoked responses in the classical conditioning task were not significantly modulated by stimulus probability (t(5) = 0.97, p = 0.81, one-sided t-test). *p<0.05. See also Fig. 3.

**Fig. S4.**
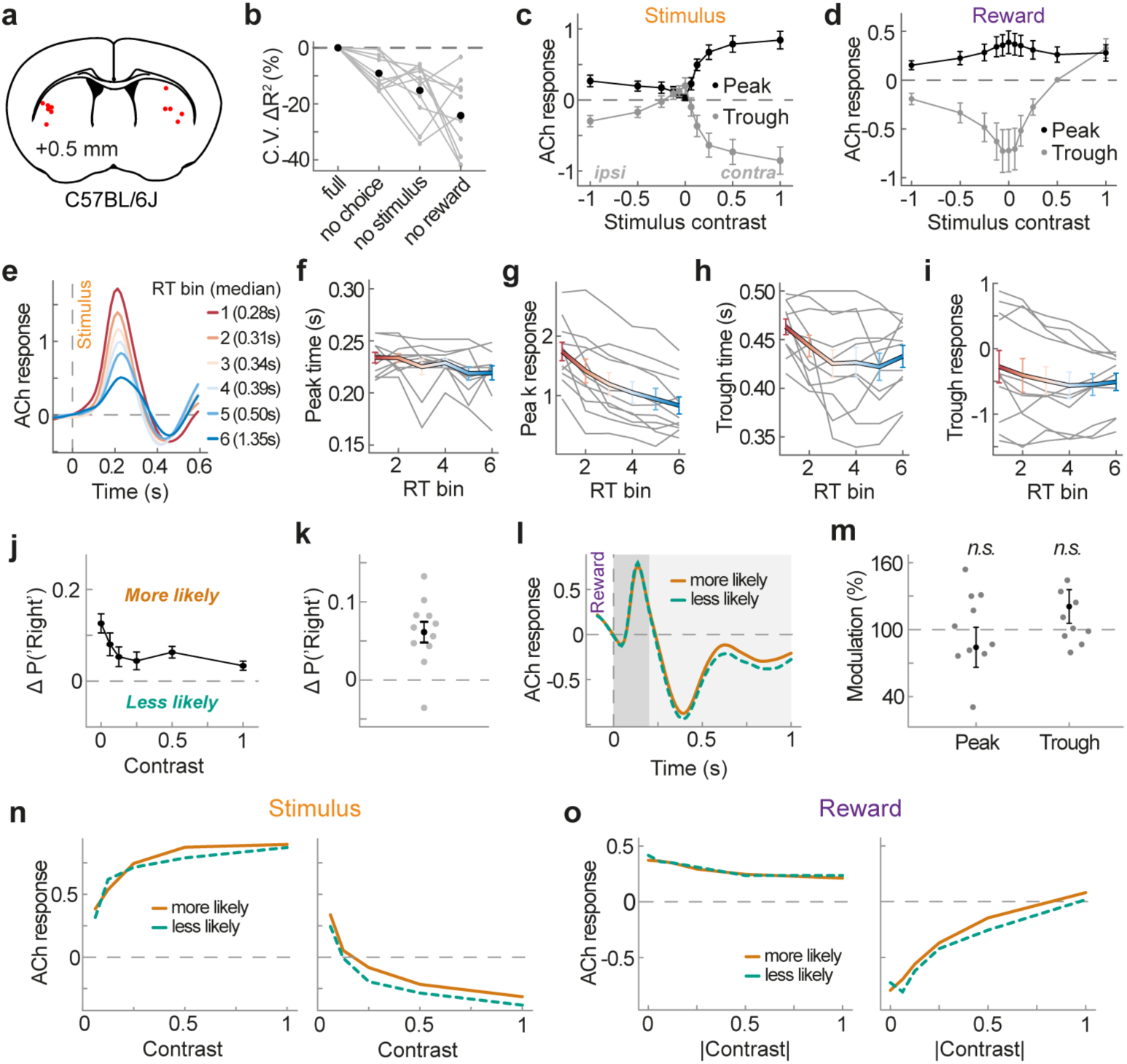
Acetylcholine recordings in the visual decision-making task. **a,** The locations of DLS optic fiber implants for acetylcholine recordings in mice performing the visual decision task with stimulus probability manipulation, estimated from post-mortem histological examinations overlaid on a brain slice schematic. **b,** Relative reduction in cross-validated R^2^ when removing choice, stimulus or reward regressors from a linear encoding model of acetylcholine release. Grey lines denote individual animals, black dots denote group averages. **c,** Group-average stimulus-evoked acetylcholine response as a function of signed stimulus contrast, relative to the recorded hemisphere. Positive x values denote contralateral stimuli. The peak (black) and trough (grey) acetylcholine response was calculated as the average response between 0.1-0.3s and 0.3-0.5s (see Fig. 4c). Error bars denote SEMs. **d,** Same as in panel c, but for reward-evoked response. Peaks and troughs were calculated between 0-0.2 s and 0.2 to 1s. **e,** Group-average stimulus-evoked acetylcholine response, split into six response time bins (red to blue), while controlling for stimulus contrast. Bin 1 contains the 1/6 fastest responses of each contrast level. **f,** Peak time of acetylcholine responses as a function of response time bin. There was a small negative relationship between response time and acetylcholine peak times (t(10) = -2.73, p = 0.02). **g,** Peak height of acetylcholine responses as a function of response time bin. Acetylcholine responses decreased with increasing response times (t(10) = -5.93, p = 0.0001). **h,** same as panel f, for trough times. There was a small negative relationship between response time and acetylcholine trough times (t(10) = -2.37, p = 0.039). **i,** same as panel g but for trough depth. There was no significant relationship between the depth of acetylcholine troughs and response times (t(10) = -1.40, p = 0.19). **j,** Difference between behavioral choice probabilities conditioned on the current high probability side, as a function of absolute stimulus contrast. Positive y values indicate a tendency to choose the high probability side. Data points show group averages. Error bars denote SEMs. **k,** Choice bias towards the higher probability side, averaged across all stimulus contrasts. Grey dots denote individual mice. Black dot shows group average. Error bars denote SEM. **i,** Group-average reward-evoked acetylcholine response, separately for high (orange) and low probability stimuli (green). **m,** Modulation of reward-evoked peak and trough acetylcholine responses by stimulus probability. Neither peak nor trough were significantly modulated (peak: 84.22 +- 17.92% of low probability response, t(9) = 0.62, p = 0.55; trough: 120.72 +- 15.81%, t(10) = 1.34, p = 0.21). Grey dots show individual mice. Black dots show group averages. Error bars denote SEMs. **n,** Group-average stimulus-evoked peak (left) and trough (right) acetylcholine responses for more (orange) or less likely stimuli (green), as a function of contralateral stimulus contrast. **o,** Group-average reward-evoked peak (left) and trough (right) acetylcholine responses for more (orange) or less likely stimuli (green), as a function of absolute stimulus contrast. See also Fig. 4.

**Fig. S5.**
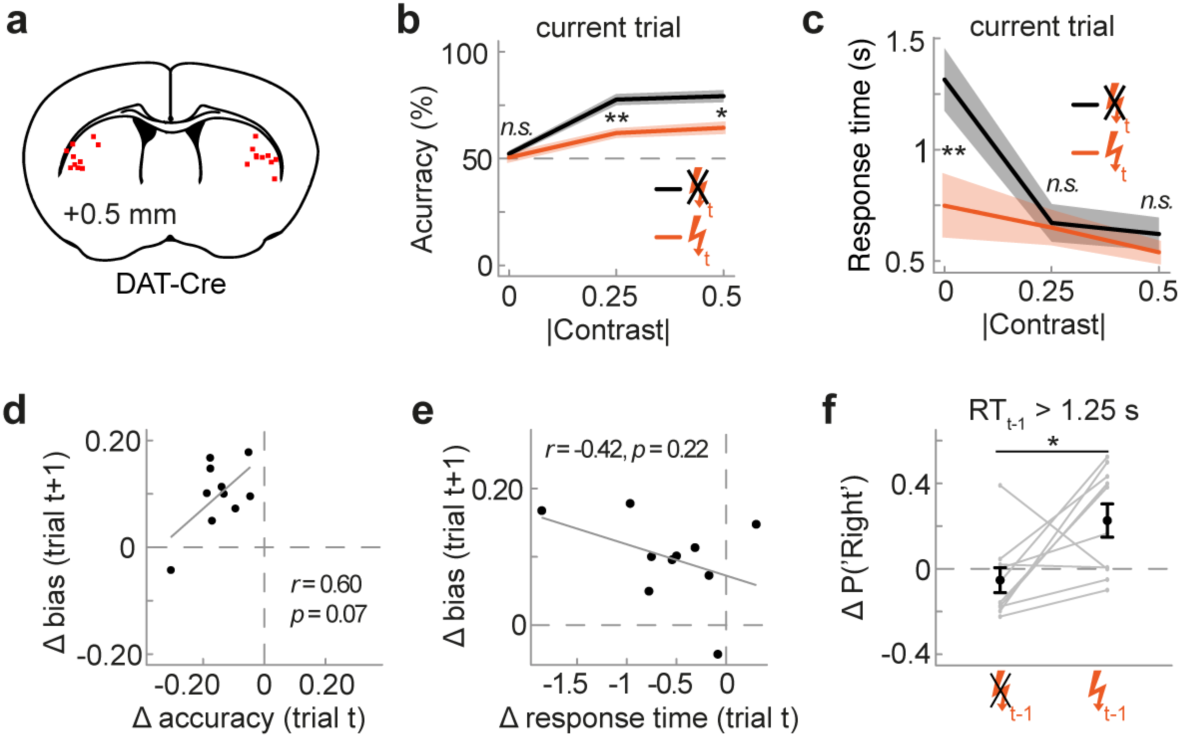
Optogenetic dopamine stimulation during visual decision-making. **a,** The locations of DLS optic fiber implants for optogenetic stimulation of dopamine release, estimated from post-mortem histological examinations overlaid on a brain slice schematic. **b,** Behavioral choice accuracy as a function of absolute stimulus contrast (x-axis), separately for stimuli with (orange) and without (black) concurrent optogenetic stimulation. Choice accuracy was decreased on stimulated relative to non-stimulated trials with 25 and 50% contrast stimuli (25%: t(9) = 2.81, p = 0.008; 50%: t(9) = 2.32, p = 0.026). Lines show group means. Shaded regions denote SEMs. **c,** Median response times as a function of absolute stimulus contrast (x-axis), separately for stimuli with (orange) and without (black) concurrent optogenetic stimulation. Response times were reduced on stimulated trials, specifically in the 0% contrast (no stimulus) condition (t(9) = 2.92, p = 0.009). **d,** Correlation between the immediate influence of optogenetic stimulation on current trial choice accuracy (x-axis) and the next trial influence on choice bias (y-axis). If next trial effects were due to current trial effects one would expect a negative correlation - larger impairments in current choice accuracy would lead to larger next trial choice biases. There was no significant correlation between the two effects, and if anything, a trend towards a positive correlation. **e,** Correlation between current trial response time and next trial choice bias effects. There was no significant correlation between the two effects. **f,** Difference between choice probabilities conditioned on the previous choice (right minus left), averaged across all contrast levels. Positive y values denote a tendency to repeat the previous choice. This analysis only considers trials in which the previous response time was longer than 1.25 seconds, i.e. where choices occurred more than 1 second after the offset of optogenetic stimulation. Similar to analyzing all trials (Fig. 5f), animals show a higher tendency to repeat the previous choice, when the previous stimulus was paired with dopamine stimulation (t(9) = -2.60, p = 0.028). Grey dots denote individual animals, black dots show group means. Error bars denote SEMs. See also Fig. 5.

**Fig. S6.**
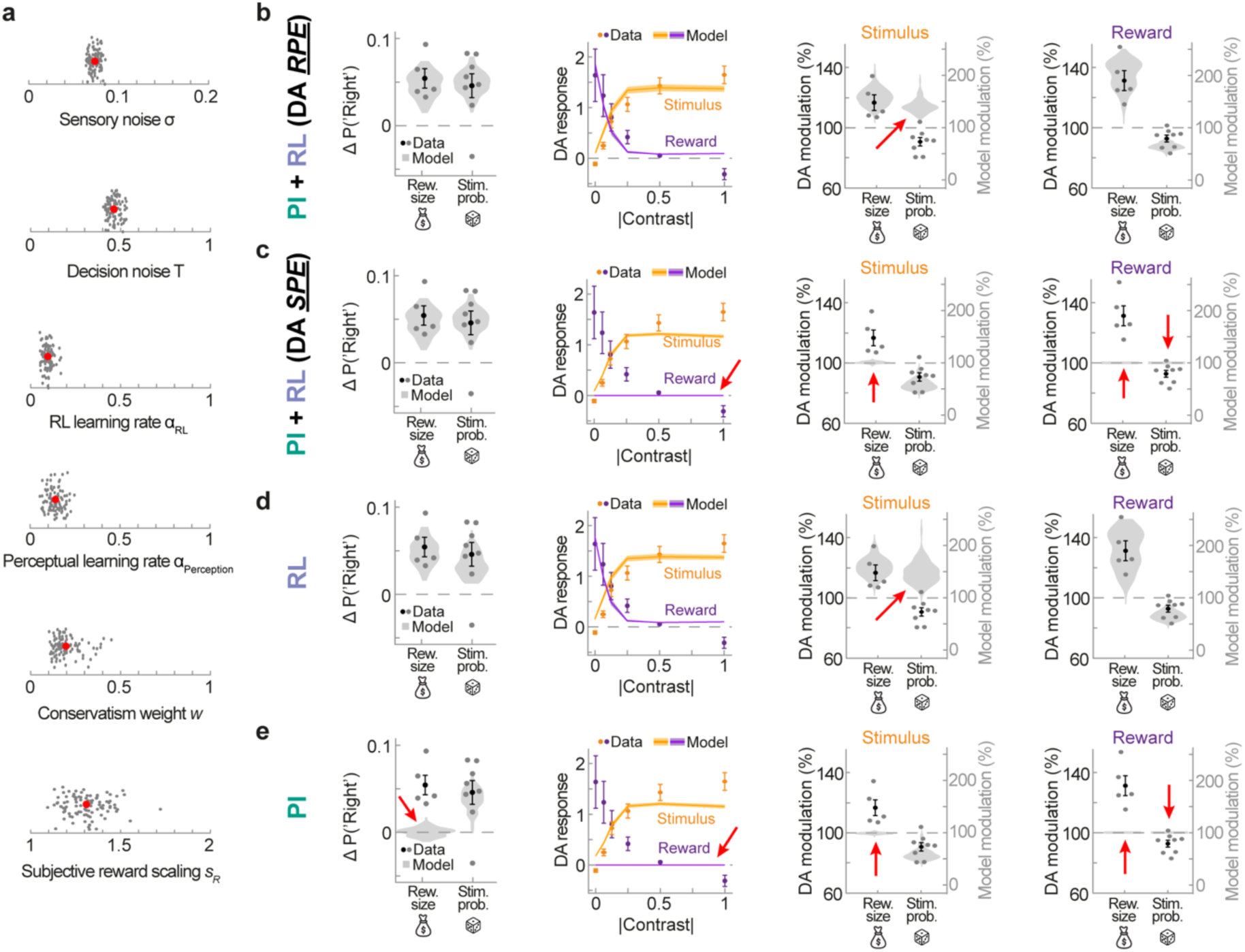
Computational models of visual decision-making behavior and dopamine release. **a,** Best fitting parameters of the PI + RL model. Grey dots denote individual fits to 100 bootstrapped data sets, resampling data of mice with replacement. Red dots denote the average parameter estimate across the bootstrap fits. **b,** Behavioral bias and dopamine signals of the PI + RL models, where dopamine encodes RPEs only. Grey and black data points show empirical mouse and group-average data. Violin plots show model estimates from 100 bootstrapped datasets. Dopaminergic RPEs fail to account for the increase in stimulus-evoked dopamine for low vs. high probability stimuli (red arrow). **c,** Same as in b, but dopamine only encodes SPEs. This model fails to account for responses in the reward size manipulation (red arrows). **d,** Behavioral bias and dopamine signals of a model engaging in reinforcement learning only. The model’s dopamine encodes RPEs, and fails to account for the increase in stimulus-evoked dopamine for low vs. high probability stimuli (red arrow). **e,** Behavioral bias and dopamine signals of a model engaging in perceptual inference only. This model fails to account for behavioral biases and dopamine signals in the reward size manipulation (red arrows). See also Fig. 6.

